# Rap1 acts via multiple mechanisms to position Canoe/Afadin and adherens junctions and mediate apical-basal polarity establishment

**DOI:** 10.1101/170977

**Authors:** Teresa T. Bonello, Kia Z. Perez-Vale, Kaelyn D. Sumigray, Mark Peifer

## Abstract

Epithelial apical-basal polarity drives assembly and function of most animal tissues. Polarity initiation requires cell-cell adherens junction assembly at the apical-basolateral boundary. The mechanisms underlying this remain key issues. Drosophila embryos provide a superb model, as 6000 polarized cells assemble simultaneously. Current data place the actin-junctional linker Canoe (mammalian Afadin’s homolog) at the top of the polarity hierarchy, where it directs Bazooka/Par3 and adherens junctions positioning. Here we define mechanisms regulating Canoe localization/function. Spatial organization of Canoe is multifaceted, involving membrane-localization, recruitment to nascent junctions and macromolecular assembly into cables at tricellular junctions. Our data suggest apical activation of the small GTPase Rap1 regulates all three events, but support multiple modes of regulation. The Rap1GEF Dizzy/PDZ-GEF is critical for Canoe tricellular junction enrichment but not apical retention. Canoe’s Rap1-interacting RA-domains mediate adherens junction and tricellular junction recruitment but are dispensable for membrane-localization. Our data also support a role for Canoe-multimerization. These multifactorial inputs all shape Canoe localization, correct Bazooka/Par3 and adherens junction positioning, and thus apical-basal polarity. We integrate existing data into a new polarity establishment model.

**Abbreviations used:** α-cat, alpha-catenin; β-cat, beta-catenin; AJ, adherens junction; Arm, Armadillo; Baz, Bazooka; CA, constitutively active; Cno, Canoe; DE-cad, *Drosophila* E-cadherin; Dzy, Dizzy; GAP, GTPase activating protein; GDP, guanosine diphosphate; GEF, guanine nucleotide exchange factor; GFP, green fluorescent protein; GTP, guanosine triphosphate; IF, immunofluorescence; MIP, maximum intensity projection; RA, Ras-associated; RFP, red fluorescent protein; SAJ, spot adherens junction; shRNA, short hairpin RNA; TCJ, tricellular junction; WT, wildtype

## Introduction

Cell polarity, the ability to differentially target proteins to distinct plasma membrane domains, is a fundamental cellular property, underlying processes from bacterial motility to neuronal transmission. We explore polarity in epithelia, the most common tissue architecture in animals, underlying organs from skin to gut to blood vessels. In epithelia apical-basal polarity distinguishes the upper (apical) and lower (basal) surfaces of each epithelial sheet, allowing them to serve as barriers between body compartments and to selectively transport molecules across this barrier(Campanale et al., 2017).

Cadherin-based cell-cell adherens junctions (AJs) reside at the junction between the apical and basolateral domains, separating them. Cadherins cell-cell mediate adhesion while proteins bound to their cytoplasmic tails, broadly known as catenins, interact with the actomyosin cytoskeleton (Meng and Takeichi, 2009). Blocking cadherin-catenin function disrupts cell adhesion and epithelial polarization in cultured cells and developing embryos (Cox et al., 1996; Gumbiner et al., 1988; Johnson et al., 1986). These advances opened up a new question: how are cadherin-catenin complexes positioned at the apical/basolateral boundary during polarity establishment?

Early *Drosophila* development provides an outstanding model of polarity establishment (Harris, 2012). Flies begin development as a syncytium, in which nuclear division occurs without cytokinesis. Nuclei move to the egg cortex and undergo several more rounds of synchronous division. They then exit the cell cycle and undergo cellularization, during which the actomyosin cytoskeleton pulls in membrane around each nucleus, creating 6000 polarized cells. The original egg cortex becomes the apical membrane, and AJs are positioned in a polarized way at the apicolateral interface.

In the absence of AJ proteins, embryos cellularize but cells then lose adhesion for one another and concurrently lose apical-basal polarity (Cox et al., 1996; Harris and Peifer, 2004). While AJs are key for polarity initiation, they themselves must be positioned apically as membranes invaginate. The polarity protein Bazooka (Baz; fly Par3) plays a key role. It colocalizes with cadherin-catenin complexes as polarity is established, in large multiprotein complexes called spot AJs (SAJs; Harris and Peifer, 2004; McGill et al., 2009; Tepass and Hartenstein, 1994). Initial small cadherin-catenin protein clusters arise early in cellularization. Baz clusters accumulating at the apicolateral interface engage these precursory cadherin-catenin complexes (McGill et al., 2009; Harris and Peifer, 2004) leading to their robust assembly into nascent SAJs, with smaller clusters all along the lateral membrane. While Baz localizes correctly in the absence of cadherins, cadherin-catenin complexes require Baz to be apically restricted. In Baz’s absence, small apical cadherin-catenin complexes localize all along the basolateral axis and fail to assemble into larger SAJs (McGill et al., 2009). This placed Baz at the top of the polarity hierarchy, opening the question of how Baz is positioned. One clue came from the fact that syncytial nuclear divisions involve polarized actin and microtubules. Strikingly, apical Baz positioning requires both dynein-based microtubule transport and an intact actin cytoskeleton (Harris and Peifer, 2005). However, the protein(s) linking nascent SAJs to the cytoskeleton remained unclear.

In the conventional model, cadherins link to actin via α- and β-catenin. Recent work revealed this is mediated by a far more sophisticated set of molecules, facilitating tension-sensing feedback mechanisms (Lecuit and Yap, 2015). The actin-binding protein Canoe (Cno; mammalian orthologue Afadin) is part of the molecular machinery linking AJs to the cytoskeleton (Mandai et al., 2013). Cno’s large multi-domain structure allows it to directly interact with the cytoskeleton via its F-actin-binding domain and bind AJ proteins including nectin, E-cadherin and α-catenin via its PDZ and proline-rich domains.

While *Drosophila cno* is not essential for cell-cell adhesion (Sawyer et al., 2009), unlike cadherin/catenins, it is required for many morphogenic processes driven by AJ/cytoskeletal linkage. Mesoderm apical construction during gastrulation offers a good example. In Cno’s absence, AJs lose connection to the contractile apical actomyosin cytoskeleton, hampering cell shape changes and mesoderm internalization (Sawyer et al., 2009). Cno plays similar roles in other actomyosin-driven processes, including germband convergent elongation and dorsal closure, helping link force-generating myosin cables to AJs (Boettner et al., 2003; Boettner and Van Aelst, 2007; Choi et al., 2011; Sawyer et al., 2011).

Afadin plays similar roles. *Afadin* null mice are embryonic lethal, with highly-disorganized ectoderm and impaired mesoderm migration (Ikeda et al., 1999; Zhadanov et al., 1999). Thus, Afadin also is not essential for cell-cell adhesion but regulates morphogenesis. Tissue-specific knockouts implicated Afadin in morphogenic events depending on cadherin function, including synaptogenesis (Beaudoin et al., 2012), lymphangiogenesis (Majima et al., 2013) and nephron lumen formation (Yang et al., 2013). In the intestine Afadin is required for normal epithelial barrier function (Tanaka-Okamoto et al., 2011) and for maintaining adhesion between Paneth and intestinal crypt cells (Tanaka-Okamoto et al., 2014).

The diverse events in which Cno/Afadin are involved and their roles in linking AJs to actomyosin suggested Cno might mediate AJ/cytoskeletal interactions during polarity establishment in cellularization. Consistent with this, Cno localizes to nascent SAJs as they form, with special enrichment in cable-like structures at tricellular junctions (TCJs), where three forming cells meet (Choi et al., 2013). Strikingly, in *cno* maternal/zygotic mutants, apical enrichment of both Baz and cadherin-catenin complexes is lost—they continue to localize to the cortex but are not apically enriched. This placed Cno at the top of the polarity establishment hierarchy, and thus raised questions about the cues localizing Cno to nascent SAJs as polarity is established.

An intact actin cytoskeleton is critical for Cno’s cortical localization (Sawyer et al., 2009). Cno localization to nascent SAJs is also dependent on the small GTPase Rap1 (Choi et al., 2013; Sawyer et al., 2009). Cno binds active Rap1 via two N-terminal Ras-associated (RA-) domains (Boettner et al., 2003). *Rap1* maternal/zygotic mutants also fail to apically enrich Baz and cadherin-catenin (Choi et al., 2013). Thus, Rap1 is an essential cue in establishing polarity, acting upstream of Cno. Consistent with a role for Rap1 in mediating AJ organization, in wing disc epithelia Rap1 helps maintain uniform AJ distribution around the cell periphery (Knox and Brown, 2002). Rap1 also helps regulate vertebrate cadherin-mediated cell adhesion (Asuri et al., 2008; Price et al., 2004; Sato et al., 2006).

Like other small GTPases, Rap1 is pleiotropic. While Rap1’s effectors vary between organisms and tissues, many act through the actin cytoskeleton (reviewed in (Frische and Zwartkruis, 2010). Drosophila *Rap1* is required for many events depending on cell shape change and dynamic junctional remodeling (Asha et al., 1999). *Rap1* loss-of-function mutants share with *cno* defects in mesoderm invagination (Sawyer et al., 2009; Spahn et al., 2012) and dorsal closure (Boettner et al., 2003; Boettner and Van Aelst, 2007). In both events, Dizzy (Dzy; fly PDZ-GEF) is the GEF regulating Rap1 (Boettner and Van Aelst, 2007; Spahn et al., 2012). The mechanisms by which Rap1 regulates Cno function and apical-basal polarization are thus areas of intense interest. Here we use the early Drosophila embryo to define these mechanisms.

## Results

### Cno recruitment to nascent SAJs and macromolecular assembly at TCJs is Rap1-dependent but Rap1-independent recruitment mechanisms begin at gastrulation

Our goal was to define mechanisms mediating apical-basal polarity establishment, with a focus on how Rap1 regulates Cno, the most upstream known player in the process. During the earliest stages of cellularization, endogenous Cno puncta are positioned apically, and move basally with the extending membrane (Fig. 1A,F). They then begin to assemble into nascent spot AJs (SAJs). Cno is initially primarily enriched at bicellular contacts (Fig. 1B,G blue arrows), colocalizing with DE-cadherin and Baz (Sawyer et al., 2009). By mid-cellularization, Cno puncta are spatially excluded from the apical domain and begin to show substantial tricellular junction (TCJ) enrichment, where Cno extends more basally (Fig. 1C,H red arrows). This enrichment and extension basally at TCJs intensifies as cellularization is completed (Fig 1D,E,I, J red arrows). Three-dimensional reconstruction revealed that at TCJs Cno forms cable-like structures extending from TCJs basally several microns (Fig. 1K,arrow).

**Figure 1.**
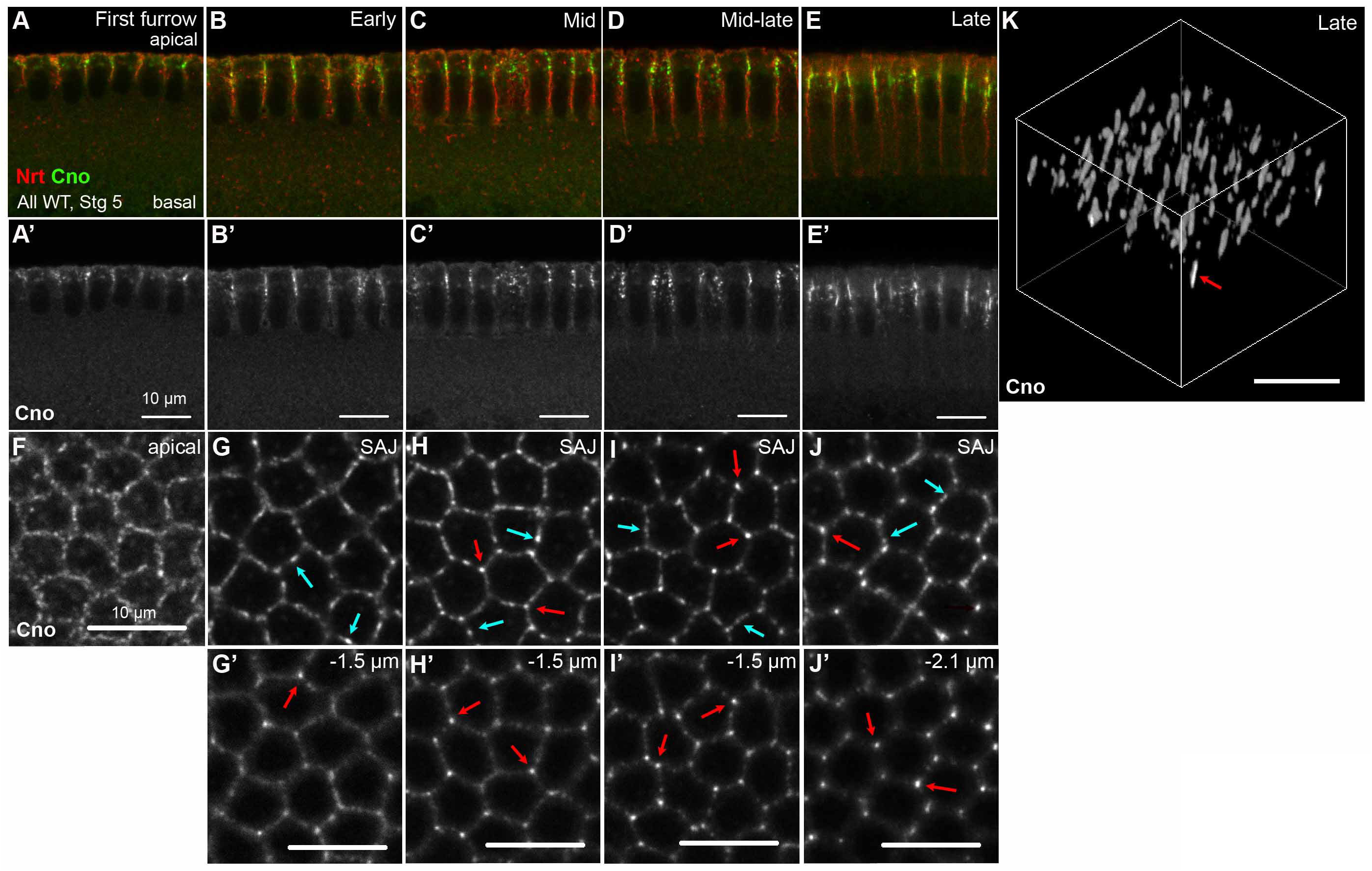
Cno localization during polarization involves apical retention at SAJs and cable formation at TCJs. (A-K) Cellularizing embryos. (A-E) Cross sections. (F-J) *En face.* (K) 3D-reconstruction. Small apical Cno puncta (A,F) move basally with the growing membrane, and are initially enriched at bicellular SAJs (B,G blue arrows). As cellularization proceeds, Cno is retained apically and becomes progressively enriched at TCJs (C-E,H-J red arrows), where it assembles into macromolecular cables (K).

Cno recruitment/maintenance at developing SAJs is Rap1-dependent. In *Rap1* maternal/zygotic mutants Cno membrane accumulation is lost, resulting in loss of apical-basal polarity and ultimate defects in epithelial integrity (Choi et al. 2013). We utilized Valium20 shRNAs targeting *Rap1* expressed under control of the GAL4-UAS system. Expression in the female germline knocks down maternal mRNA, and maternally-expressed GAL4 persists zygotically, driving zygotic shRNA expression, often leading to knockdown mimicking maternal/zygotic mutants (Staller et al., 2013). This approach reduced Rap1 below levels detectable by immunoblotting (Suppl. Fig. 1G,H), effectively reproducing the *Rap1* null phenotype (Suppl. Fig. 2A,B). In late cellularizing *Rap1* RNAi embryos, Cno is virtually absent from the membrane in apical-basal cross sections (Fig. 2A,B), maximum intensity projections (MIPs) of cross-sections (Fig. 2C,D) or apical *en face* sections (Fig. 2I,J; all images intensity matched with wildtype). Reduced Cno at the membrane is a consequence of altered localization and not a change in absolute Cno levels (Suppl. Fig. 1G).

**Figure 2.**
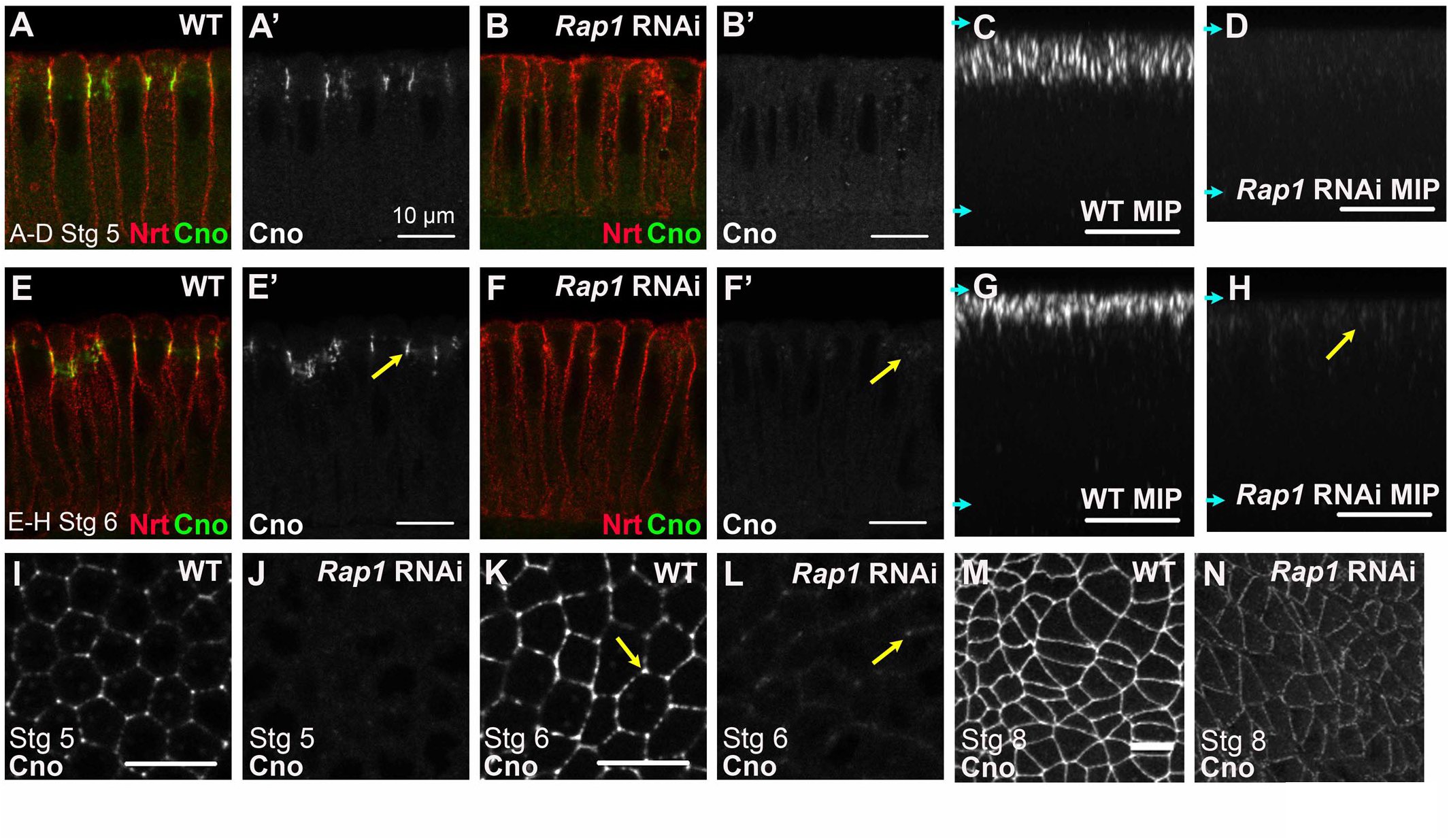
Cno localization during polarity establishment is Rap1-dependent but localization is restored as gastrulation proceeds. Intensity-matched images, wildtype (WT) vs. *Rap1* RNAi. (A,B,E,F) Cross sections. (C,D,G,H) Maximum intensity projections of cross sections (MIPs). (I-N) *En face.* (A-D,I,J) Late stage 5. Rap1 knockdown largely eliminates Cno localization to the membrane. (A vs B, single sections; C vs D; MIPs; I vs J, *En face* views). (E-H,K,L) Late stage 6. Initial weak reappearance of Cno at apical SAJs and TCJs (arrows). (M,N) Cno AJ localization is partially restored by germband extension (apical 2.2 µm MIP).

The polarity regime established during cellularization is elaborated on during gastrulation. At this point, new polarity cues come into place, and, for example, apical Baz and AJ localization is partially restored in *cno* mutants (Choi et al., 2013). We thus examined whether Cno localization was restored in *Rap1* RNAi embryos when gastrulation began. Low-intensity Cno puncta reappeared by late stage 6, which are apically enriched (Fig. 2E-H,arrows) and associated with TCJs (Fig. 2K,L,arrows), but overall Cno intensity at the membrane remained very low. Thus, Rap1 still plays a role during early polarity maintenance. However, as germband extension neared completion, Cno localization to AJs resumed (Fig. 2M,N), though at reduced levels. Thus, Cno localization is strongly Rap1-dependent during polarity establishment but becomes more Rap1-independent in the polarity maintenance phase. This Cno relocalization is not sufficient to maintain ectodermal integrity, however, as *Rap1* RNAi embryos exhibit strong epithelial disruption (Suppl. Fig. 2B).

### Active Rap1 is necessary and sufficient to recruit Cno to the cellularizing membrane

Rap1 localization during cellularization contrasts with Cno’s polarized localization at the apicolateral boundary. Endogenous Rap1 is not polarized (Choi et al., 2013), instead localizing along the entire length of the cellularizing membrane, extending down to the furrow front where actin and myosin are enriched. Rap1’s physical interaction with Cno/Afadin is dependent on Rap1’s activity state (Boettner et al., 2000; Boettner et al., 2003). Since Rap1 itself is not polarized, we hypothesized Rap1 activation is spatially restricted to the apicolateral boundary and consequently defines the site for initial Cno localization and SAJ formation.

To manipulate Rap1 <u>activity</u> we created three GAL4-UAS-driven GFP-tagged Rap1 transgenes(Ellis et al., 2013): Rap1WT, Rap1S17A, which is locked in the GDP-bound off state (referred to as Rap1GDP), and the constitutively GTP-bound and thus active Rap1Q63E (referred to as Rap1CA). Small GTPase mutants like Rap1S17A trap GEFs in high-affinity complexes, effectively sequestering them from acting on downstream GTPases (Dupuy et al., 2005). Conversely, constitutively-active mutants are locked in the GTP-bound state and rendered insensitive to GAPs, resulting in continuous signaling to downstream effectors (Kitayama et al., 1990; Scrima et al., 2008). When expressed using maternal-GAL4, Rap1WT, Rap1CA and Rap1GDP were 11, 15 and 3.5-fold above endogenous Rap1 levels, respectively (1–4 h, Suppl. Fig. 3A,B). All three localized along the entire lateral membrane during cellularization (Fig. 3A-C), like endogenous Rap1 (Choi et al., 2013). Expressing Rap1WT had no effect on embryo viability (Suppl. Fig. 2E). In contrast, expressing either GDP-locked or GTP-locked Rap1 mutants was embryonic lethal (Suppl. Fig. 2E). Rap1GDP strongly disrupted epithelial integrity, with the ventral epidermis fragmented (Suppl. Fig. 2C,F), similar to, though slightly more severe than, *Rap1* maternal/zygotic loss (Choi et al., 2013) or RNAi (Suppl. Fig. 2B). In contrast, Rap1CA embryos had a largely intact epidermis, with defects in head involution alone or in conjunction with epidermal holes (Suppl. Fig 2D,F).

**Figure 3.**
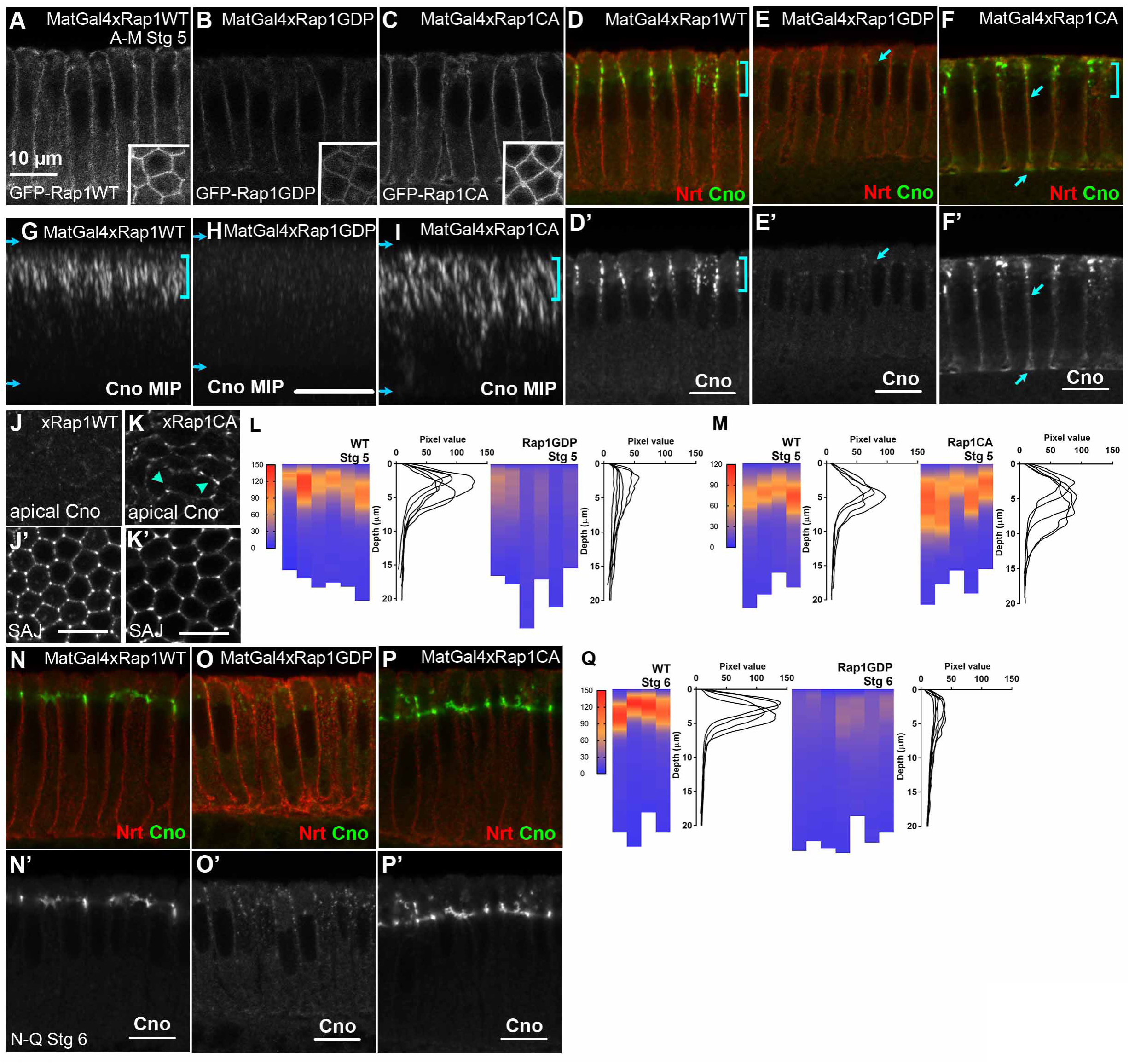
Active Rap1 is necessary and sufficient to recruit Cno to the cellularizing membrane. (A-M) Cellularization=late stage 5. (A-C) GFP-tagged Rap1WT, Rap1GDP, and Rap1CA all localize uniformly along the membrane as viewed in lateral and *en face* (insets) sections. (D,G) In embryos expressing Rap1WT, Cno localizes normally to the nascent SAJs at cellularization (brackets). (E,H,L). In embryos expressing Rap1GDP, Cno localization to the membrane is drastically reduced. (F,I,M). In embryos expressing Rap1CA, Cno puncta expand both apically (J vs K, *en face* arrowheads) and basally (I vs G, MIPs) and a uniform pool of Cno extends along the basolateral axis (F, arrows). (L,M) Quantification of localization changes. (N-Q) Gastrulation onset=late stage 6. (N) Rap1WT does not affect Cno localization to apically-focused AJs. (O, quantified in Q) Rap1GDP. Small Cno puncta begin to appear along the lateral membrane but fail to effectively cluster. (P) Rap1CA. Cno focuses into AJs but not as tightly as in wildtype.

We next examined how altering Rap1 activity affects Cno localization. In embryos overexpressing Rap1WT, Cno localizes normally to nascent SAJs during cellularization (Fig. 3D,G MIP). At gastrulation onset Cno clustered at the extreme apical end of the cell, as in wildtype (Fig. 3N). In contrast, eliminating Rap1 <u>activation</u> by expressing Rap1GDP resulted in almost complete loss of Cno from the membrane (Fig. 3E, H), thus mimicking *Rap1* null or RNAi embryos (Fig. 2; Choi et al., 2013). Small Cno puncta were observed along the length of the invaginating membrane, but their low-level intensity suggests they do not account for all Cno displaced from nascent SAJs (Fig. 3E,H). We quantified this using ImageJ PlotProfile to measure pixel-intensity in projected cross sections compared to intensity matched controls (Fig. 3L, as in Choi et al., 2013). When the polarity maintenance phase initiates, Cno levels at the membrane were not restored in Rap1GDP embryos (Fig. 3O; measured in Q). However, as in *Rap1* RNAi embryos, low-intensity Cno puncta appeared on the membrane by late stage 6 (Fig. 3O’).

Our hypothesis that a spatially-confined pool of active Rap1 is an essential cue for Cno polarization predicted expressing Rap1CA along the length of the cellularizing membrane would broaden Cno membrane recruitment basally. Consistent with this, Rap1CA induced several changes in endogenous Cno localization. In Rap1CA embryos, a subset of membrane-associated Cno did not exhibit Cno’s characteristic apically-restricted, punctate appearance, but rather localized in a uniform manner all along the basolateral membrane (Fig. 3F, arrows). This is reminiscent of Rap1CA’s localization (Fig. 3C). A second Cno pool remained apically enriched (Fig. 3F, bracket), but relative to wildtype there were two differences. First, the Cno puncta in Rap1CA embryos extended more basally, involving, in part, basal extension of TCJ Cno cables (Fig. 3G vs I; quantified in M). Second, Cno puncta were found more apically in Rap1CA embryos (Fig. 3J vs K arrowheads; *en face* views at SAJs vs more apical). As Rap1CA embryos gastrulate, mislocalized Cno puncta are excluded from the apical domain and become more tightly focused at the AJs (Fig. 3P). Interestingly however, apical AJ translocation in the dorsal ectoderm is delayed in Rap1CA (Fig. 3N vs P), thus exaggerating the normal difference in timing of wildtype AJ translocation between dorsal and ventral ectoderm (Weng and Wieschaus, 2016). Together, these data are consistent with the idea that an early apical asymmetric distribution of active Rap1 along the cellularizing membrane regulates Cno’s recruitment to nascent SAJs.

### Loss of Rap1 <u>activity</u> mimics total Rap1 loss, disrupting Baz and AJ polarization

Cno and Rap1 regulate polarity establishment by regulating apical Baz positioning. In the absence of either Cno or Rap1, Baz puncta lose apical restriction and redistribute all along basolateral membrane (Choi et al., 2013). While expressing Rap1WT did not affect Baz polarization (Fig. 4B,F vs A,E), Rap1GDP expression depolarized Baz. Membrane-associated Baz puncta extended all along the basolateral membrane and up into the apical domain above the nascent SAJs (Fig. 4C,G vs A,E). This phenotype was partially rescued as embryos entered gastrulation; Baz was gradually excluded from the lateral domain, accompanied by enrichment at the apicolateral interface (Fig. 4L vs K). Expressing Rap1GDP also disrupted cadherin-catenin complex assembly into SAJs (Fig. 5A vs B), thus resembling *Rap1* or *cno* null embryos (Choi et al., 2013). The pool of smaller Arm-puncta along the basolateral membrane and enrichment at basal junctions (Fig. 5A,B arrows) appeared unaffected. AJ protein polarization was not substantially restored at gastrulation onset (Fig. 5F,I vs E,H). Thus, Rap1 activity is required to initiate Baz and AJ polarization and plays a role in AJ polarity maintenance.

**Figure 4.**
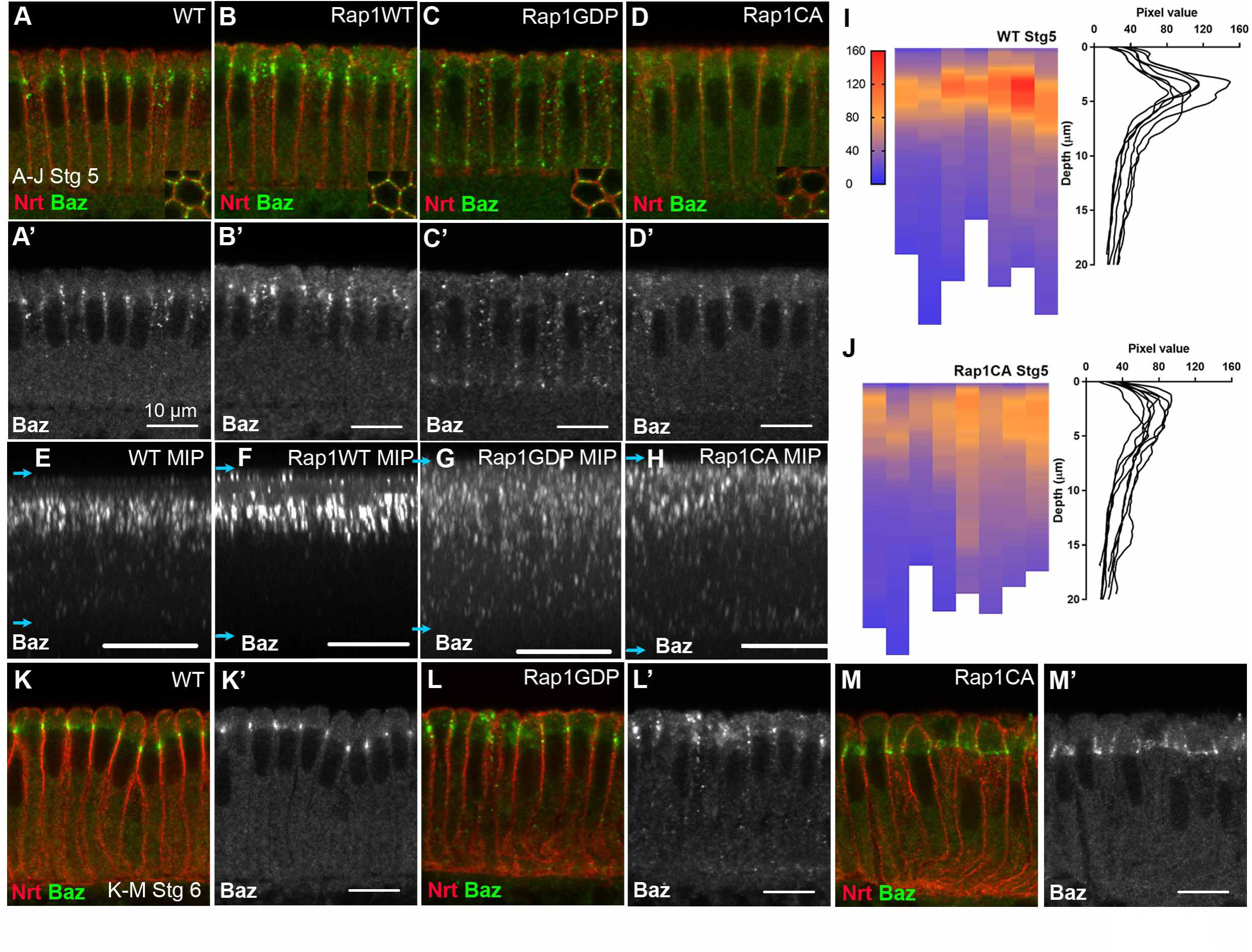
Manipulating Rap1 activity perturbs Baz polarization. (A-J) Cellularization=late stage 5. A-D insets= *en face* views at SAJ level. (A,E) Baz is restricted to the apicolateral SAJs. (B,F) Expressing Rap1WT does not perturb Baz polarity. (C,G) Baz polarization is lost in embryos expressing Rap1GDP. Small Baz puncta are redistributed along the basolateral axis, but remain membrane-associated (C, inset). (D,H,I vs J) Expressing Rap1CA disrupts the tight restriction of Baz to forming SAJs-it localizes both more apically and more basally, reducing Baz peak intensity at the apicolateral membrane. (K-M) Late stage 6. (K) In wildtype Baz puncta undergo enhanced apically clustering. (L) In Rap1GDP embryos, Baz puncta are cleared from the basolateral domain but fail to effectively cluster. (M) In Rap1CA embryos, Baz is cleared from the basolateral domain and exhibits clustering, albeit at reduced levels.

**Figure 5.**
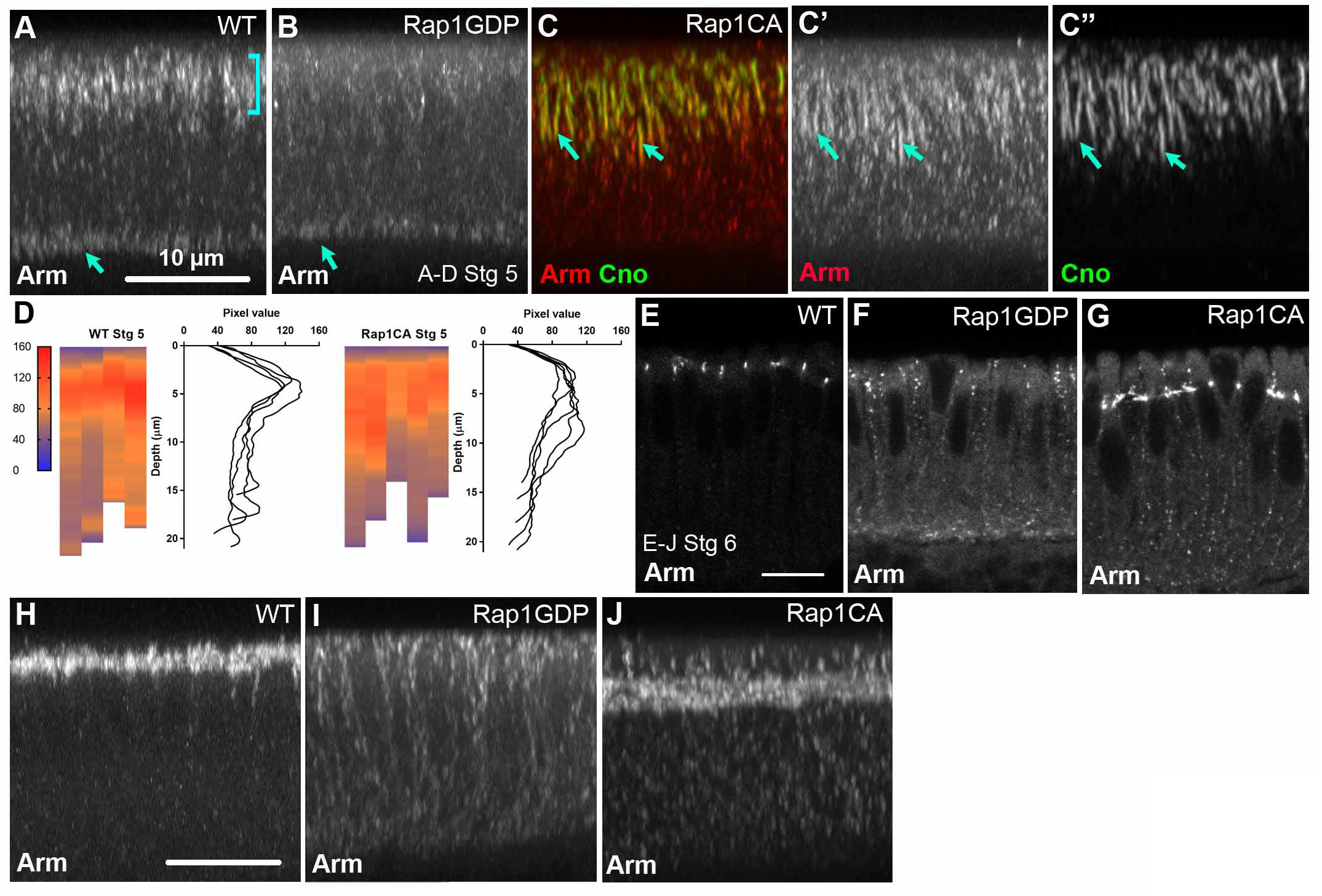
Polarized Rap1 activity is required for apical AJ positioning. (A-D) Cellularization=late stage 5. Maximum intensity projections (MIPs). (A) Wildtype. Arm localizes to apical SAJs (bracket) and basal junctions (arrow). (B) Rap1GDP. Arm no longer enriched at SAJs. (C) Rap1CA. Arm expands basally and localizes to Cno cables (arrows). (D) Quantification. Arm is less focused at SAJs in Rap1CA. (E-J) Gastrulation onset=Late stage 6. (F,I vs E,H) Rap1GDP. Little recovery of Arm in AJs. (G,J) Rap1CA. Polarized Arm clustering occurs but apical translocation of AJs in the dorsal ectoderm is delayed.

### Unrestricted Rap1 activity reduces apical Baz and AJ enrichment

We next examined how unrestricted Rap1 activity affected Baz polarization. In Rap1CA embryos, Baz puncta became enriched in the apical half of the forming cell but did not concentrate as tightly at nascent SAJs (Fig. 4D,H vs A,E). Pixel intensity plots confirmed this flattening of the Baz peak relative to wildtype (Fig. 4I vs J). Apical Baz clustering was largely restored at gastrulation onset (Fig. 4M vs K), though focusing of Baz into AJs was less complete. In cellularizing Rap1CA embryos, AJ proteins remained somewhat enriched apically but were not as tightly apically restricted, localizing along the upper half of the lateral membrane in a manner similar to Baz (Fig. 5A vs C’, quantified in D). Mislocalized Arm puncta largely co-localized with the extended Cno cables (Fig. 5C,arrows), suggesting that while Rap1CA disrupts precise spatial organization of AJs, molecular interactions between AJ proteins remain at least partially intact. Arm colocalization with Cno did not extend to the smoothly localized pool of Cno along the basolateral domain; instead Arm localized to small clusters as in wildtype. AJs assembly began to be restored at gastrulation, though apical translocation was delayed (Fig. 5G,J). These data suggest apical Cno clustering and subsequent Baz and Arm recruitment to nascent AJs are regulated by localized Rap1 activity. Thus, localized Rap1 activity is necessary for establishing polarity, but is less essential for maintaining the polarized state.

### The Rap1 GEF Dizzy specifically regulates Cno enrichment at TCJs

One mechanism to spatially compartmentalize Rap1 activity is via specific GEFs. The Rap1GEF Dzy (fly PDZ-GEF) is a plausible candidate, given its role in mesoderm invagination, which immediately precedes cellularization and requires Rap1 and Cno (Spahn et al., 2012; Sawyer et al., 2009). We generated embryos lacking maternal/zygotic Dzy, crossing females with germlines homozygous for the null allele *dzy^Δ1^* (Huelsmann et al., 2006) to *dzy^Δ1^* heterozygous fathers. 64% of progeny died as embryos, suggesting partial zygotic rescue of the 50% of embryos receiving paternal wildtype *dzy*. Dead embryos had defects in head involution, dorsal closure and epidermal integrity (Suppl. Fig. 2G,H). We then examined effects on Cno localization. In contrast to *Rap1* mutants (Choi et al., 2013), Cno was not lost from the plasma membrane. Cno was apically restricted to nascent SAJs (Fig. 6B,D vs A,C; localized to the apical 50±3% of the cell in *dzy* (n=7) vs. 45±6% of the cell in wildtype (n=3)). However, there was one striking difference from wildtype. At mid-late cellularization, wildtype embryos exhibited strong Cno enrichment at TCJs, both at the level of SAJs (Fig. 6C) and 2µm basally (Fig. 6C’). In contrast, in *dzy* embryos Cno localized to bicellular contacts but was not enriched at TCJs (7/7 late stage 5 embryos, Fig. 6D vs C). This defective assembly of TCJ Cno cables was even more obvious in 3-dimensional reconstructions (Fig. 6E,F). The TCJ defect persisted into gastrulation (3/5 stage 6 embryos, Fig. 6G,J vs H,K), but zygotic rescue was observed in ∼50% of embryos (2/5 embryos, Fig. 6I,L). Thus, Dzy is not essential for initial Cno membrane-recruitment or apical-basal positioning, but is required for effective TCJ clustering. This suggests Rap1 is regulated by more than one cue, with different regulatory inputs coordinating Rap1’s role in Cno polarization versus junctional maturation. *dzy* mutants also exhibited more irregular apical cell shapes during cellularization (Fig. 6M vs N), a phenotype of *Rap1* but not *cno* mutants (Choi et al., 2013), consistent with Dzy also regulating this Rap1 activity. *dzy* mutants also mimicked effects of *cno* (Choi et al., 2011; Sawyer et al., 2011; Sawyer et al., 2009) in later events, disrupting mesoderm invagination (Spahn et al., 2012), exaggerating Baz planar polarization during germband extension and leading to deepened segmental grooves during dorsal closure (Suppl. Fig 2I-P). These data suggest Dzy is an important regulator of Rap1 and Cno during many morphogenetic events, but is not the only regulator.

**Figure 6.**
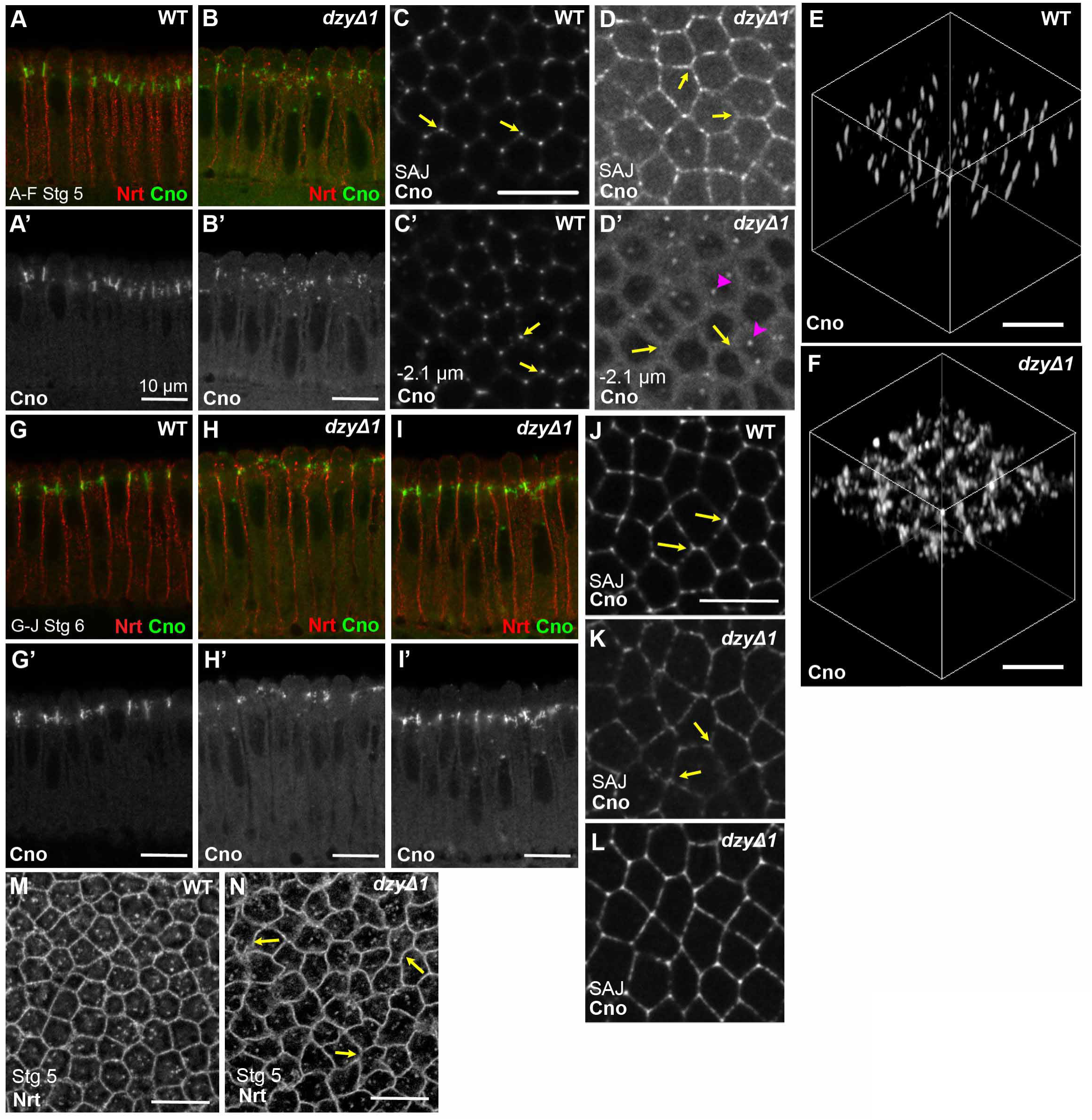
The GEF Dizzy is required for Cno enrichment at TCJs. (A,B,G-I) Cross sections. (C,D,J-N) *En face.* (E,F) 3D-reconstructions. (A vs B) Cno puncta continue to localize to membrane and are apically restricted in *dzy* mutants. (C vs D) In *dzy* mutants Cno remains at SAJs but is not enriched at TCJs (arrows) and does not extend basally there. Some enrichment at centrosomes is observed (arrowheads). (G-I) Cno localization at gastrulation onset in wildtype (G,I), putative maternal/zygotic (H,K) or zygotically-rescued (I,L) *dzy* mutants. TCJ enrichment remains reduced in maternal/zygotic mutants (arrows). (M,N) Irregular apical cell shapes (arrows) in cellularizing *dzy* mutants.

### Cno’s RA-domains are dispensable for Cno membrane recruitment but required for apical retention and tricellular junction clustering

Cno’s RA-domains bind constitutively-active Rap1 (Boettner et al., 2003). To define the RA-domain’s roles in Cno regulation and function, we created three C-terminally GFP-tagged Cno constructs driven by the GAL4-UAS system: full-length wildtype Cno (CnoWT), Cno with the N-terminal RA-domains cleanly deleted (Cno∆RA), or Cno with the F-actin-binding domain and linker region removed in addition to the RA-domains (CnoFHA-PDZ). When driven by maternal GAL4, all accumulated at similar levels (Suppl. Fig. 1A,B,D). Overexpressing CnoWT or CnoΔRA in a wildtype background did not appreciably affect embryo lethality (data not shown).

One simple mechanism by which Rap1 could recruit Cno to the plasma membrane and nascent SAJs is via direct interaction with the RA-domains. In the simplest version of this hypothesis, deleting the RA-domains would have the same effect on Cno as removing Rap1, abolishing membrane recruitment. To test this hypothesis, we wanted to express Cno∆RA in a background in which endogenous Cno was reduced/eliminated. We were fortunate that an existing shRNA targeted the 5’-region of *cno* mRNA deleted in Cno∆RA. This allowed us to knockdown endogenous Cno without affecting Cno∆RA. Endogenous Cno was effectively knocked down during cellularization/gastrulation (Suppl. Fig. 1E,F, <5%). Cno continued to be substantially depleted during mid-late embryogenesis, though there was some rebound in expression.

We thus expressed either CnoWT-GFP or Cno∆RA-GFP in embryos with endogenous Cno knocked down by RNAi. The shRNA also targeted CnoWT-GFP. However, fortuitously, the high-level expression of CnoWT-GFP partially balanced this, reducing its levels to near those of endogenous Cno in controls (Suppl. Fig. 1E,F). The GFP-signal was sufficiently strong to confirm that CnoWT-GFP retained apical enrichment, localizing like endogenous Cno in wildtype (Fig. 7C,F vs A,D).

**Figure 7.**
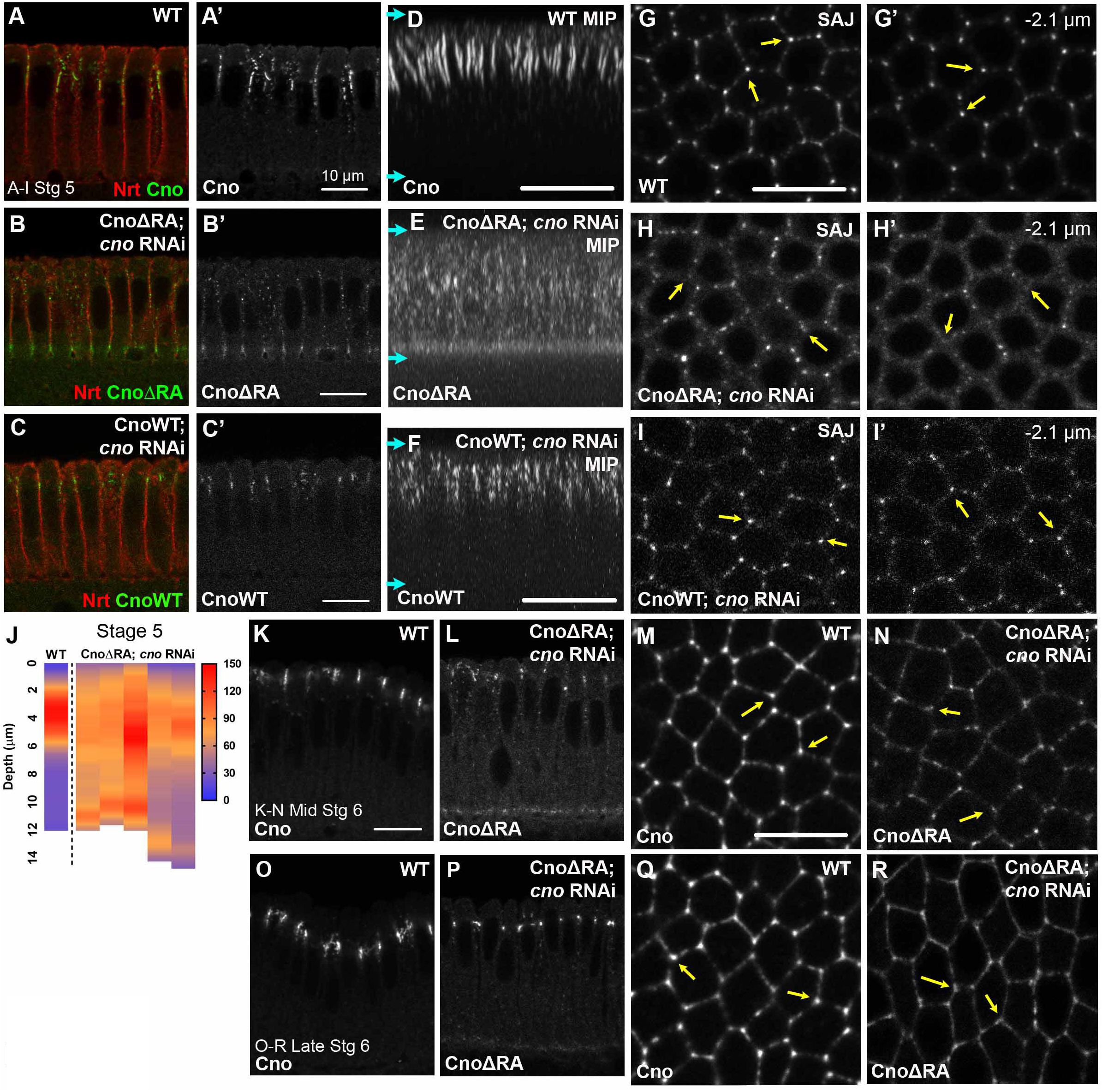
Cno’s RA-domains are required for effective apical retention and TCJ clustering. CnoWT::GFP or Cno∆RA::GFP expressed in *cno* RNAi background, visualized with GFP-tag. (A-C,K,L,O,P) Cross sections. (D-F) Maximum intensity projections (MIPs). (G-I,M,N,Q,R) *En face.* (A-J) Cellularization. Cno∆RA (B,E) is less tightly restricted apically than endogenous Cno (A,D) or CnoWT (C,I; since CnoWT is also knocked down by the shRNA, its levels are lower than endogenous Cno or Cno∆RA). Cno∆RA (H,H’) is also less reliably enriched in TCJs (arrows) than endogenous Cno (G,G’) or CnoWT (I,I’). (J) Quantification, Cno∆RA apical restriction. (K-R) Gastrulation onset=mid-late stage 6. Apical enrichment of CnoΔRA::GFP is partially-rescued by mid-stage 6 (K vs L) and strongly rescued by late-stage 6 (O vs P), but TCJ enrichment remains reduced (M vs N, Q vs R arrows).

This knockdown-rescue approach allowed us to test whether Cno∆RA was deficient in either membrane recruitment or apical enrichment. Strikingly, CnoΔRA expressed in a *cno* RNAi background remained membrane-associated (Fig. 7B,H vs. A,G), in contrast to Cno in *Rap1* mutants. However, in contrast to CnoWT expressed in a *cno* RNAi background (Fig 7F), Cno∆RA apical enrichment was severely reduced (Fig. 7B,E; quantified in J). Furthermore, CnoΔRA was not particularly enriched at TCJs versus neighboring bicellular junctions (Fig. 7H vs. G,arrows), unlike CnoWT (Fig. 7I). At gastrulation onset, Cno∆RA began to be recruited into apical AJs (Fig. 7L), but apical clustering and TCJ enrichment remained impaired (mid-stage 6; Fig. 7L,N vs K,M). By late stage 6 Cno∆RA re-localized to AJs but not TCJs (Fig. 7O-R). These data suggest Cno∆RA can localize to the membrane independently of endogenous Cno, but it is defective in apical retention at SAJs and clustering at TCJs. This is consistent with the idea that different inputs regulate different aspects of Cno localization.

### Rap1 is essential for Cno∆RA recruitment to SAJs but both CnoWT and Cno∆RA are less reliant on Rap1 for localization after gastrulation onset

In parallel, we considered an alternative mechanism by which Rap1 could regulate Cno localization. Actin-regulatory formins provide an interesting model. Their N-terminal RA-domain folds back onto the rest of the protein, locking it into a closed, inactive conformation. This is relieved by Rho binding(Rose et al., 2005). We thus tested the hypothesis that Cno’s RA-domains are similarly auto-inhibitory, with inhibition relieved by Rap1 binding. If so, deleting the RA-domains might allow Rap1-independent Cno SAJ recruitment. To test this, we expressed either CnoWT-GFP or Cno∆RA-GFP in *Rap1* RNAi embryos. We confirmed knockdown by immunoblotting—during cellularization Rap1 was knocked down to levels undetectable by immunoblotting (Suppl. Fig. 1G,H). As expected, *Rap1* RNAi eliminated apical enrichment of CnoWT-GFP in nascent SAJs (Fig. 8A vs B, arrowheads), though weak localization of small puncta all along the lateral membrane remained (Fig. 8F, arrows). CnoWT-GFP accumulated ectopically at basal junctions (Fig. 8B,D arrows), something we also saw weakly when expressing CnoWT in a wildtype background. Strikingly, *Rap1* RNAi had similar effects on Cno∆RA-GFP localization, eliminating apical enrichment at SAJs (Fig. 8C, arrowhead) and leading to ectopic basal junction accumulation (Fig. 8C,E arrows). There was little rescue of either CnoWT or Cno∆RA localization to AJs at gastrulation onset (stage 6; Fig. 8J,K). Thus deleting the RA-domains does not simply open up an inactive conformation of Cno to render it Rap1-independent for localization.

**Figure 8.**
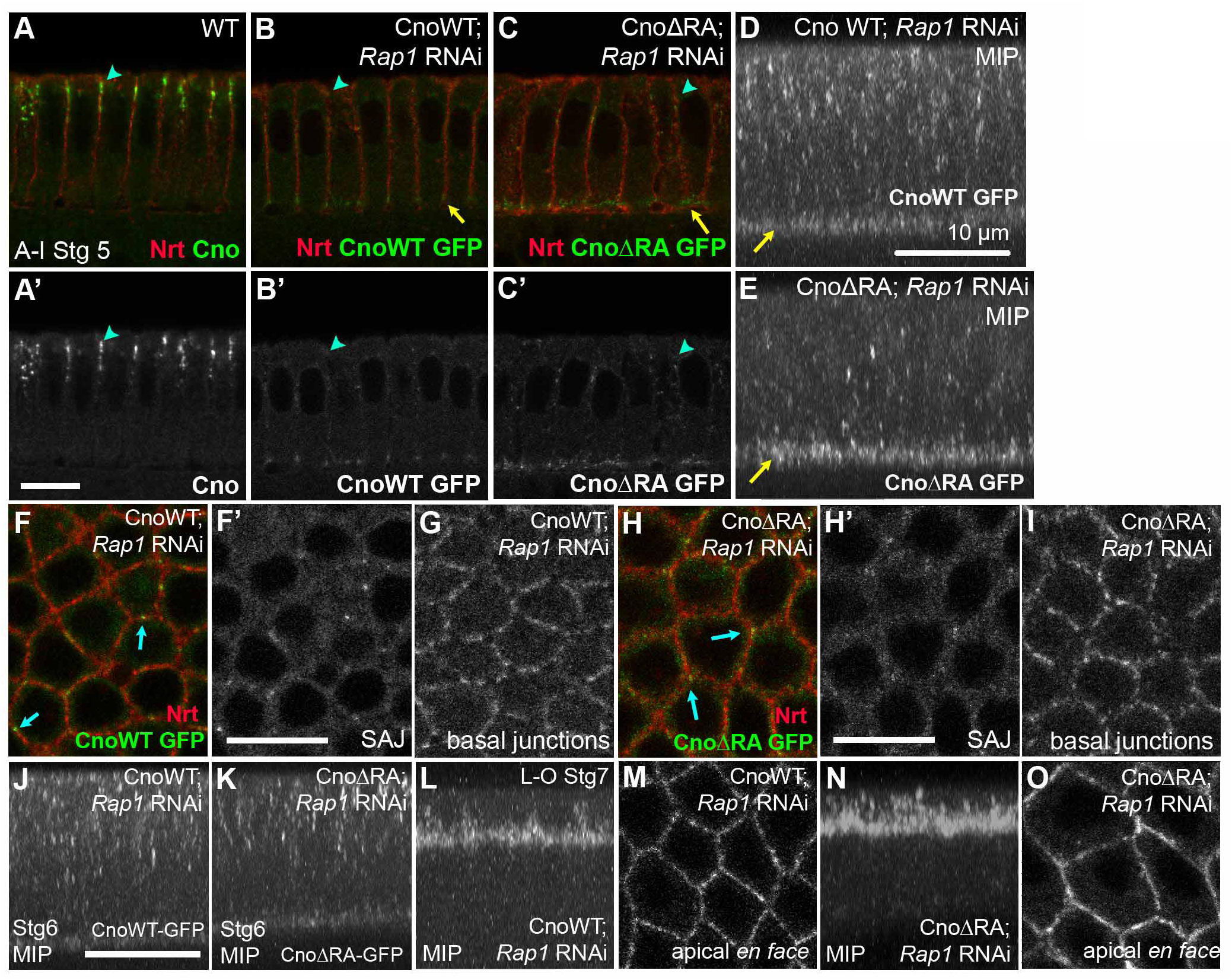
Rap1 is essential for cortical recruitment of Cno∆RA. CnoWT::GFP or CnoΔRA::GFP expressed in *Rap1* RNAi background, visualized with GFP-tag. (A-I) Stage 5. Both CnoWT::GFP (B,D,F,G) and CnoΔRA::GFP (C,E,H,I) are lost from SAJs in Rap1’s absence (arrowheads)--the few remaining puncta are membrane-associated. (F,H). For both, mislocalized Cno remains at basal junctions (B-E arrows, G,I). (J,K) MIPs, stage 6. Mislocalization persists. (L-O) Stage 7. CnoWT::GFP and CnoΔRA::GFP begin to relocalize to AJs.

After gastrulation begins, initial defects in AJ protein localization in some mutants begin to be rescued, restoring normal or near-normal junctional localization—Par1 activation can contribute to this (McKinley and Harris, 2012). After Rap1 knockdown endogenous Cno relocalizes to AJs as gastrulation proceeds (Fig. 2; Rap1 knockdown remained quite effective even at 12–15hr; Suppl. Fig. 1G,H). Similarly, when germband extension initiated (stage 7) both CnoWT and Cno∆RA relocalized to AJs in *Rap1* RNAi embryos (Fig. 8L-O). Thus Cno is less reliant on Rap1 for post-gastrulation localization.

### Endogenous wildtype Cno rescues Cno∆RA’s localization defects

One striking aspect of Cno’s localization is its assembly into macromolecular “cables” at TCJs. This suggested it might have the ability to self-interact. We thus tested whether this putative interaction would allow wildtype Cno to recruit Cno∆RA to nascent SAJs. To test this, we expressed our GFP-tagged Cno constructs in a wildtype background, using maternal-GAL4. CnoWT-GFP largely replicated endogenous Cno localization. It was enriched at nascent SAJs as they formed (Fig 9A,B), with particular enrichment in TCJ cables (Fig. 9C,D). There also was slight enrichment at the basal junctions, just apical to the actomyosin machinery driving membrane invagination—we suspect this is an over-expression artifact. We next examined Cno∆RA-GFP. In contrast to Cno∆RA localization defects after *cno* RNAi, in a wildtype background Cno∆RA localization also strongly resembled that of endogenous Cno, with strong recruitment to forming apical SAJs (Fig. 9E,F) and marked enrichment at TCJs, where it assembled into cables (Fig. 9G,H). Like CnoWT-GFP, there also was some basal junction enrichment. Both CnoWT and Cno∆RA also resembled endogenous Cno in becoming more focused in the apical-basal axis in late stage 6 (Fig. 9I,J) and moving apically with AJs at stage 7 (Fig. 9K,L). Thus, in the presence of wildtype endogenous Cno and Rap1, the RA-domain is dispensable for Cno localization. In contrast, deleting the F-actin binding domain and linker in addition to the RA-domains (CnoFHA-PDZ), completely abolished membrane localization (Fig. 9M). This suggests that the F-actin binding or linker regions are important for membrane localization.

**Figure 9.**
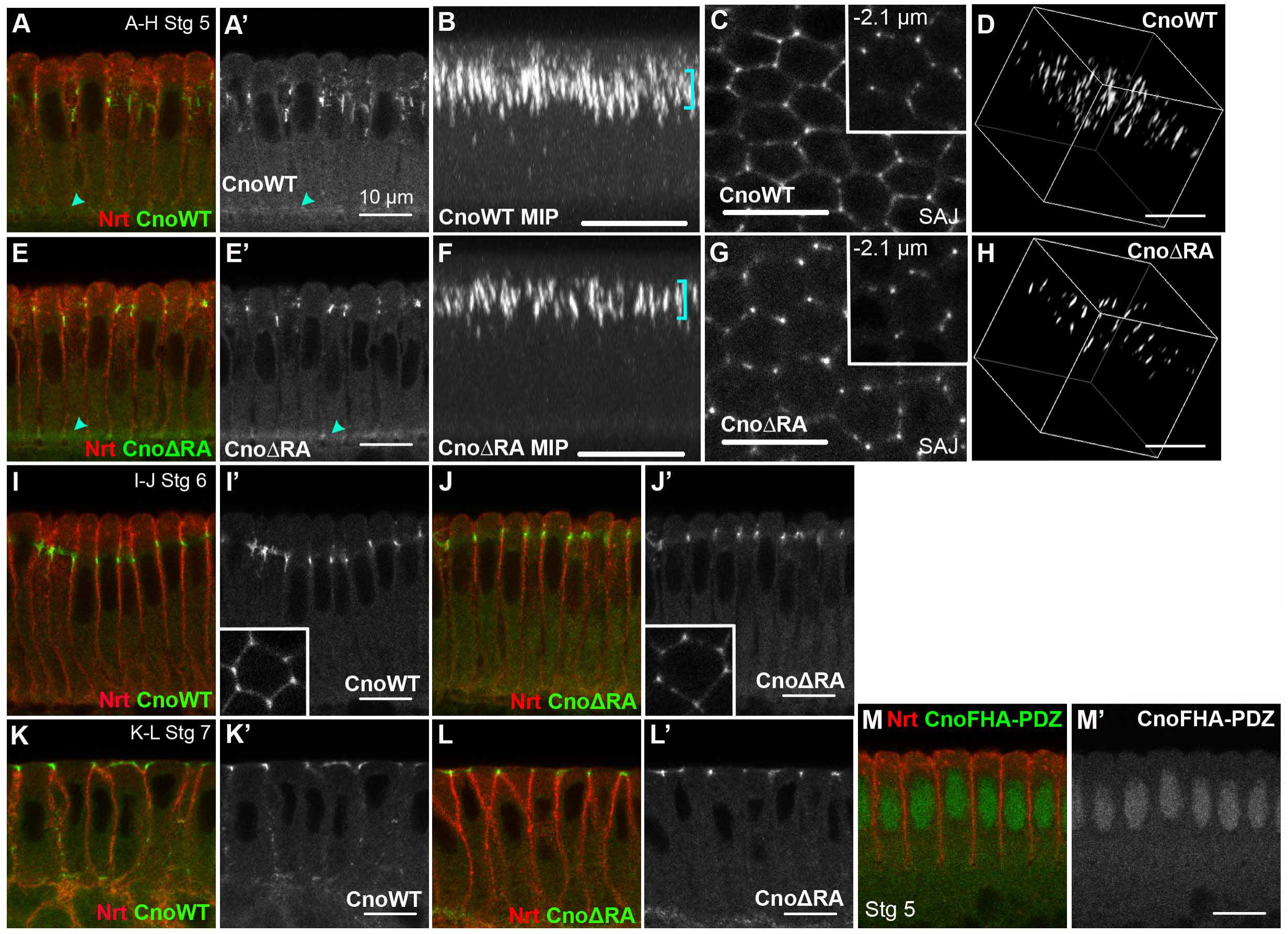
Endogenous Cno rescues CnoΔRA’s localization defects. CnoWT::GFP or CnoΔRA::GFP expressed in wildtype background, visualized with GFP-tag. (A-H) Cellularization=stage 5. CnoWT::GFP (A,B) and CnoΔRA::GFP (E,F) localize to nascent SAJs (brackets) and basal junctions (arrowheads). (C,G) *En face*. Both CnoWT::GFP (C) and CnoΔRA::GFP (G) proteins are enriched at TCJs at the level of SAJs and 2.1µm basally (insets). (D,H) 3-dimensional projections. Both CnoWT::GFP (D) and CnoΔRA::GFP (H) localize to cables at TCJs. (I,J) Late stage 6. CnoWT::GFP (I) and CnoΔRA::GFP (J) undergo apicolateral clustering, like endogenous Cno. (K,L) Stage 7. CnoWT::GFP and CnoΔRA::GFP move apically, like endogenous Cno. (M) CnoFHA-PDZ::GFP does not localize to the membrane.

### Cno∆RA retains significant though not full function

In our final tests, we examined Cno∆RA function. We first examined its ability to rescue *cno* RNAi knockdown, which leads to highly-penetrant embryonic lethality (Fig. 10I) and strong defects in morphogenesis (Fig. 10A-E, J). Since combining two UAS-driven transgenes can reduce expression, as a control we expressed both *cno* RNAi and red fluorescent protein (RFP), controlling for the possibility that Cno knockdown was reduced. We observed no significant reduction in Cno knockdown (Suppl. Fig. 1E,F) or embryonic lethality (Fig. 10I), although effects on morphogenesis were slightly alleviated (Fig. 10J). Expressing CnoWT in *cno* RNAi embryos completely rescued both embryonic viability (Fig. 10I) and morphogenesis, as assessed by cuticle phenotype (Fig. 10A-E,J) and embryo staining (Fig. 10F,G). Cno∆RA also retained substantial function, though less than CnoWT, despite accumulating at much higher levels. Embryonic lethality was reduced to 41% (Fig. 10I), and in dead embryos morphogenesis was essentially fully restored (Fig. 10J,H).

**Figure 10.**
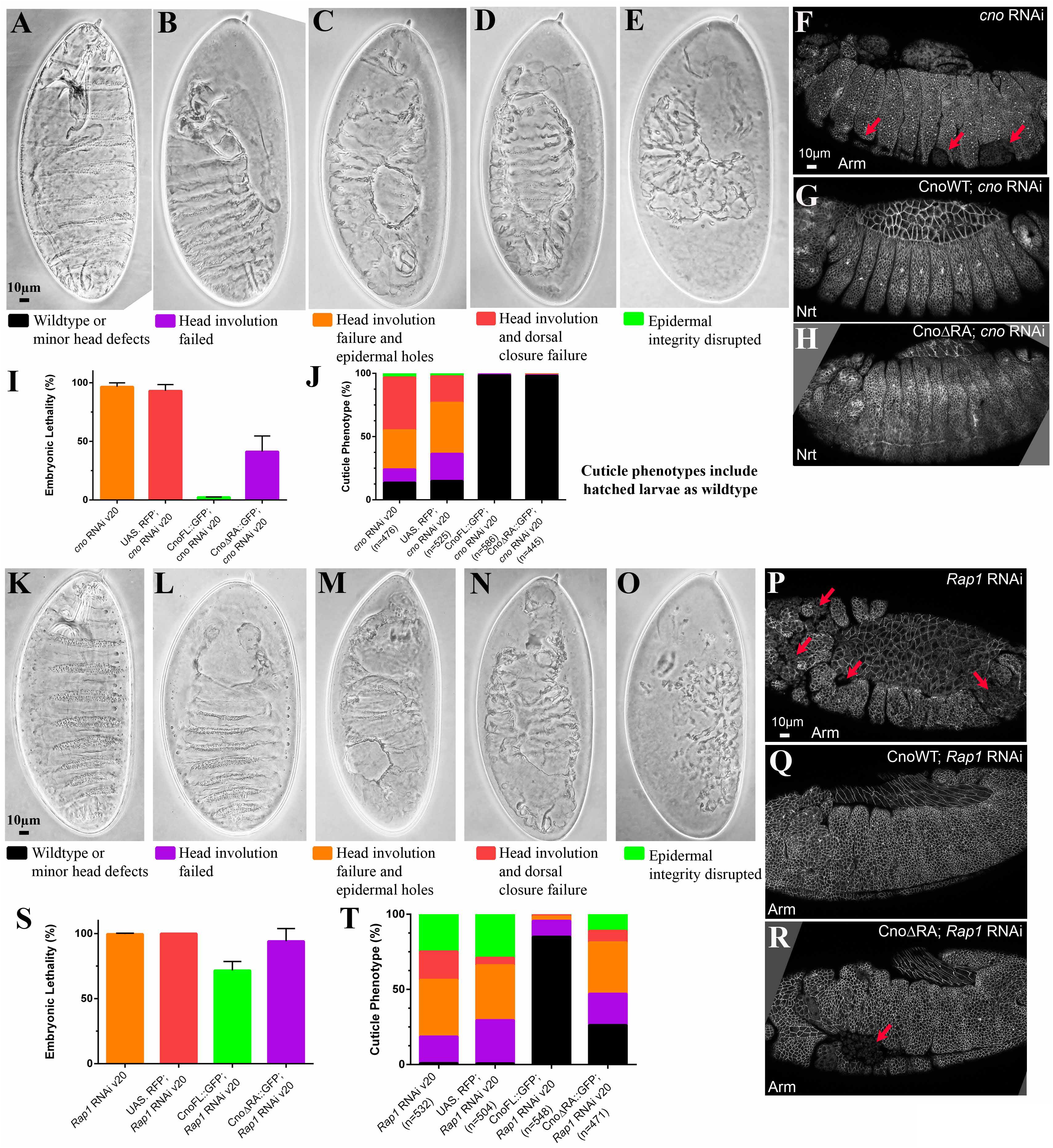
Cno∆RA retains significant though not full function. (A-E,K-O) Range of morphogenesis defects in cuticles of *cno* RNAi (A-E) or *Rap1* RNAi embryos (K-O) with or without expression of CnoWT or Cno∆RA. (F-H,P-R) Representative antibody-stained embryos. (I,S) Effects on embryonic lethality of expressing maternally-driven CnoWT or Cno∆RA in *cno* RNAi (I) or *Rap1* RNAi (S) backgrounds. (J,U) Quantitation, effects on morphogenesis and cuticle phenotype of expressing maternally-driven CnoWT or Cno∆RA in *cno* RNAi (J) or *Rap1* RNAi (T) backgrounds.

The *Rap1* RNAi background, with its severely reduced Rap1 levels, provided a sensitized background in which to compare Cno∆RA and CnoWT function. *Rap1* RNAi led to completely-penetrant embryonic lethality (Fig. 10S), and strongly disrupted head involution, dorsal closure, and, in severe cases, epidermal integrity (Fig. 10K-O,T). We once again controlled for two transgenes using RFP, with little or no effect on knockdown (Suppl. Fig. 1G,H) or phenotype (Fig. 10S,T).

We then examined whether CnoWT or Cno∆RA rescued Rap1 knockdown. CnoWT partially rescued embryonic viability (Fig. 10T). Morphogenesis rescue was more substantial; 85% of embryos had a nearly wildtype epidermis (Fig. 10Q,T). In contrast, expressing Cno∆RA in the *Rap1* RNAi background led to no rescue of embryonic viability (Fig. 10S), and the rescue of embryonic morphogenesis was significantly less complete (Fig. 10R,T). Thus Cno∆RA retains significant but not full function.

## Discussion

Apical-basal polarity establishment is a key step in animal development, driving development of epithelial tissues and organs and creating the architecture enabling many morphogenetic movements. Early fly embryos provide a premier model of this process. Work from many labs defined assembly and apical positioning of cell-cell AJs as the key initial step. Efforts then focused on defining molecular and cellular mechanisms underlying this, revealing key roles for Baz/Par3 (Harris and Peifer, 2004), and more recently, the junctional-actin crosslinker Cno and its upstream activator Rap1 (Choi et al., 2013). This re-focused our attention on the next level of mechanistic analysis: how does Rap1 regulate Cno localization and do Rap1-independent cues also play a role?

### Apical Rap1 activity promotes Cno junctional recruitment at the top of the polarity hierarchy

Our current model of apical-basal polarity establishment during fly embryogenesis suggests the key upstream step is positioning Cno at the site where AJs will form. The small GTPase Rap1 and F-actin play roles in this process (Sawyer et al., 2009). However, the mechanism by which Rap1 acted remained unclear, as Rap1 localizes all along the invaginating membrane. Our new data support a model in which apical Rap1 activity plays a key role. The ability of GTP-locked constitutively-active Rap1 to recruit Cno all along the basolateral membrane and the loss of Cno from the membrane induced by GDP-locked Rap1 are both consistent with this. In the future, it would be valuable to design a Rap1 activity sensor to confirm this hypothesis. As we discuss below, it was intriguing that although Cno’s RA-domain plays a role in apical Cno recruitment/retention, Rap1 also influences localization of a *cno* mutant lacking this domain.

Our next task was to define Rap1 activation mechanisms. Rap1 has many GEFs, including C3G, Epac, CalDAGGEF1 and PDZ-GEF (Gloerich and Bos, 2011). Dzy, the mammalian PDZ-GEF homolog, was the most likely candidate, as it has a role 30 minutes later in mesoderm apical constriction, which also requires Rap1 and Cno (Spahn et al., 2012). Strikingly, although our analyses suggest Dzy regulates Cno in later morphogenetic events like germband extension, junctional planar polarization, and segmental groove retraction, maternal/zygotic Dzy loss did not fully mimic effects of Rap1 loss on polarity establishment. Dzy loss replicated only a subset of these effects—cortical Cno recruitment and apical restriction were unaffected. Instead, Dzy loss specifically affected Cno assembly into multiprotein cables at TCJs, and also led to defects in columnar cell shape like those caused by Rap1 loss. Based on this, we hypothesize that multiple GEFs regulate Rap1 activity, each directing specific aspects of cellularization and polarity establishment. Each might create temporally- or spatially-restricted pools of active Rap1, with different Rap1 pools mediating different effects on Cno localization. In the future, it will be important to investigate other Rap1GEFs like C3G and Sponge. Another alternative is apical restriction of Rap1 activation via basolateral Rap1GAPs, as occurs at other times (Wang et al., 2013)—this will be important to explore.

### The RA-domain plays important roles in Cno localization and activity but Cno∆RA retains significant function

Cno’s N-terminus carries two RA-domains that bind Rap1 (Boettner et al., 2003). Given Rap1’s essential role in Cno localization and function, we suspected the RA-domains would be similarly essential. We tested two hypotheses for the mechanism by which Rap1 could act via the RA-domains to regulate Cno function during polarity establishment: 1) Rap1-GTP binding to the RA-domain opens a closed, autoinhibited conformation, or 2) Rap1-GTP binding to the RA-domain physically recruits Cno to sites where AJs will be assembled. Our data essentially rule out the first hypothesis, as it predicted that Cno∆RA recruitment to nascent AJs would be Rap1-independent. Our data are largely consistent with the second hypothesis, though Rap1’s effect on Cno∆RA localization revealed that the Rap1-RA-domain interaction is not the only means of regulation.

Our data also suggest that after gastrulation onset Rap1-independent mechanisms of Cno localization come into play, as cortical localization of both wildtype Cno and Cno∆RA is restored at that stage in Rap1 knockdown embryos. While some residual Rap1 may remain, we think this is unlikely as we cannot detect it by immunoblotting at stages when Cno localization is restored. Previous work supports this, as Cno can localize to the cortex during dorsal closure in embryos expressing GDP-locked Rap1 (Boettner et al., 2003). One potential mechanism of post-gastrulation Cno recruitment is via its known interactions with AJ proteins. Before AJs assemble, Rap1 may be essential for cortical Cno recruitment but once AJs reappear after gastrulation onset, interactions between Cno and α-cat or DE-cad may restore Cno recruitment to AJs. Both CnoWT and Cno∆RA also retained significant function in Rap1 knockdown embryos, suggesting Cno has Rap1-independent activity, at least at elevated levels. Cno∆RA was significantly less effective than CnoWT, confirming and extending earlier work during dorsal closure (Boettner et al., 2003). However, Cno∆RA retained significant function in Cno knockdown embryos, suggesting the RA-domain is not absolutely essential. It may facilitate some Cno/Afadin activities: e.g., mammalian Afadin’s RA-domain regulates interactions with p120catenin and modulates E-cadherin endocytosis (Hoshino et al., 2005).

### Cno may act as a coincidence detector, with multiple simultaneous inputs regulating its positioning and that of nascent AJs

Several observations support the hypothesis that Cno localization responds to multiple inputs. Rap1 activity plays a key role, as Cno cannot localize to the membrane in Rap1’s absence or when its ability to load GTP is compromised. It seems likely that direct interactions between Rap1 and Cno’s RA-domain play a role in this. However, since Cno∆RA still requires Rap1 to localize correctly, this suggests additional complexity. A second Rap1 interaction site may exist in Cno outside the RA-domain.

Alternately, other Rap1 effectors may regulate Cno localization by different mechanisms. During cellularization, Rap1 has Cno-independent effects on apical contractility and thus columnar cell shape (Choi et al., 2013). Perhaps this postulated effector alters the actomyosin cytoskeleton in a way promoting Cno binding, since intact actin is required for Cno cortical localization (Sawyer et al., 2009). However, Cno does not simply co-localize with actin or myosin, as both are most enriched at the leading edge of the invaginating membrane. Perhaps there is an apical pool of actin in a particular conformation or with particular binding partners that allow Cno to “choose” the correct localization. Consistent with an important role for Cno’s C-terminal actin-binding domain or intrinsically disordered linker in localization, CnoFHA-PDZ did not localize cortically in wildtype embryos, unlike Cno∆RA. Together these data suggest a multi-factorial recruitment mechanism.

### A revised model of junctional assembly and polarity establishment

The Drosophila embryo affords unmatched temporal resolution, allowing us to follow AJ morphogenesis both in the context of apical-basal polarity establishment, and later in polarity maintenance. This complex process begins with the appearance of the cadherin-catenin clusters, which arise independently of a Baz polarization cue (McGill et al., 2009). It continues through formation of mature SAJs, which coordinate with the contractile cytoskeleton to enable the first morphogenetic movements of gastrulation. We now view these events as involving two independent but interlocked processes: apical restriction of Cno and AJ proteins, and their assembly into higher-order multiprotein complexes. Integrating our data with previous analyses prompts the following model. Baz helps ensure that small cadherin-catenin complexes present before cellularization are recruited/retained at an apicolateral position and assembled into larger complexes containing >1000 cadherin molecules(McGill et al., 2009). Cno plays an important role in this, helping retain Baz at the apicolateral site of SAJ assembly. Our new data suggest Rap1 activity guides two important aspects of AJ morphogenesis–AJ protein recruitment/retention at the apicolateral interface and specialized assembly of larger macromolecular complexes at TCJs. Our data further suggest these two events require at least two spatial cues directed by active Rap1, one acting via Cno’s RA-domains and one independent of that. Our results with Rap1GEF Dzy suggest these two events have different modes of regulation. Finally, our data are consistent with a Cno-driven, self-reinforcing feedback loop. Correct Cno localization can recruit more Cno to the membrane. This clustering is particularly prominent at TCJs. Intriguingly, recent work in mammalian cells suggest actomyosin cables anchor end-on at TCJs and Afadin acts there to maintain trapezoidal cell shapes (Choi et al., 2016). Whether Cno oligimerization is intrinsic to Cno itself, or is mediated by other partners, is an important question for future work. These data also have potential implications for mammalian Afadin’s roles in epithelial polarity in the kidney and intestine, another place for further research.

## Materials and Methods

### Fly stocks

Fly stocks used in this study are listed in Table 1. Mutations are described at FlyBase (http://flybase.org). Wildtype was *yellow white* or *Histone-GFP*. All experiments were done at 25°C. The stock to make *dzy* germline clones was kindly provided by Rolf Reuter (Universität Tübingen, Tübingen, Germany; Huelsmann et al., 2006). The *dzy* germline clones were made by heat shocking 48- to 72-h old *hsFLP^1^; FRT2L dzy^∆1^/FTR40Aovo^D1–18^* larvae for 3 h at 37°C (Chou et al., 1993). shRNA knockdown of *Rap1* and *cno* was carried out by crossing double copy *mat-tub-GAL4* females (Staller et al., 2013) to males carrying UAS.*Rap1* short hairpin RNA interference (shRNAi) or UAS.*cno* shRNAi (VALIUM20; Ni et al., 2011). Maternal expression of GFP-tagged Cno constructs or Rap1 activity mutants was carried out by crossing males to female double copy *mat-tub-GAL4* flies.

**Table 1:** Fly stocks, antibodies, and probes. IF, immunofluorescence. WB, western blotting.

| Fly Stocks |  | Source |
| --- | --- | --- |
| UAS.*Rap1* RNAi (stock #35047) |  | Bloomington *Drosophila* Stock Center (Bloomington, IL) |
| UAS.*cno* RNAi (stock #33367) |  | Bloomington *Drosophila* Stock Center |
| FRT2L *dzy*<sup>Δ1</sup>/Sm1 |  | R. Reuter (Universität Tübingen, Tübingen, Germany) |
| UAS.CnoWT, UAS.CnoΔRA, UAS.CnoFHA-PDZ |  | See Fly stocks in materials and methods |
| UAS.Rap1<sup>WT</sup>, UAS.Rap1<sup>S17A</sup>, UAS.Rap1<sup>Q63E</sup> |  | See Fly stocks in materials and methods |
| Antibodies/probes (species) | Dilution | Source |
| **Primary antibodies:** |  |  |
| anti-Neurotactin (mouse) | 1:100 (IF) | Developmental Studies Hybridoma Bank (University of Iowa, Iowa City, IA) |
| anti-Canoe (rabbit) | 1:1,000 (IF, WB) | J. Sawyer and N. Harris (University of North Carolina, Chapel Hill, NC) |
| anti-Bazooka (rabbit) | 1:2,000 (IF) | Choi *et al.*, 2013 |
| anti-Armadillo (mouse) | 1:50 (IF), 1:500 (WB) | Developmental Studies Hybridoma Bank |
| anti-Rap1 (rabbit) | 1:500 (WB) | Choi *et al.*, 2013 |
| anti-α-Tubulin (mouse) | 1:5,000 (WB) | Sigma Life Science (St. Louis, MO) |
| anti-Y-Tubulin (mouse) | 1:2,000 (WB) | Sigma Life Science |
| anti-GFP (mouse) | 1:1,000 (WB) | Clontech Laboratories (Mountain View, CA) |
| **Secondary antibodies:** |  |  |
| Alexa 488, 568, and 647 | 1:1,000 (IF) | Life Technologies (Carlsbad, CA) |
| IRDye 680RD, IRDye 800CW | 1:10,000 (WB) | LI-COR Biosciences (Lincoln, NE) |

### Transgenic fly lines

Rap1WT, Rap1Q63E and Rap1S17A were described previously (Ellis et al., 2013). All Rap1 activity mutants are tagged with GFP at the N-terminus. Rap1S17A is inserted on the second chromosome, Rap1WT and Rap1Q63E are inserted on the third. To generate *cno* transgenic lines, full length Drosophila *cno* cDNA was PCR-cloned into the pCR8/GW/TOPO Gateway Entry Vector (Life Technologies, Grand Island, NY). Using this template, Cno domain deletions were generated using the following forward/reverse oligonucleotide primers:

CnoΔRA, Forward: 5’CGTCCGGCGGACTCACAGCCCCGGAGGAGAAAGAAAAAA-3’

Reverse: 5’TTAGTGCACCGCGTCTATATCTCG-3’

CnoFHA-PDZ, Forward: 5’CGTCCGGCGGACTCACAGCCCCGGAGGAGAAAGAAAAAA;

Reverse: 5’TAGATGGCTCCCTGCTTGGCCACTTCC-3’

Cno mutant constructs were then recombined into Drosophila UASp Gateway vectors. CnoΔRA corresponds to deletion of amino acids 1–345. CnoFHA-PDZ corresponds to amino acids 346–1100. All Cno transgenic lines are C-terminally tagged with GFP and targeted to the second chromosome.

### Cuticle preparation

Cuticle preparation was performed according to Wieschaus and Nüsslein-Volhard (1986).

### Immunofluorescence

Antibodies used are listed in Table 1. Dechorionated embryos were fixed in boiling Triton salt solution (0.03% Triton X-100, 68 mM NaCl, 8 mM EGTA) for 10 s followed by fast cooling on ice and devitellinized by vigorous shaking in 1:1 heptane:methanol. Embryos were stored in 95% methanol/5% 0.5 M EGTA for at least 48 h at −20˚C prior to staining. Embryos were washed thrice with 0.01% Triton X-100 in PBS (PBS-T), followed by blocking in 1% normal goat serum (NGS) in PBS-T for 1 h. Primary and secondary antibody staining were both carried out at 4°C overnight with nutation. Antibody dilutions are listed in Table 1.

### Image acquisition and manipulation

Fixed embryos were mounted in Aqua-poly/Mount and imaged on a confocal laser-scanning microscope (LSM 710 or LSM 880; 40x/NA 1.3 Plan-Apochromat oil objective; Carl Zeiss, Jena Germany). ZEN 2009 software was used to process images and render z-stacks in three-dimension. Photoshop CS6 (Adobe, San Jose, CA) was used to adjust input levels so that the signal spanned the entire output grayscale and to adjust brightness and contrast. Analysis of apical-basal positioning on maximum intensity projections was performed as previously described (Choi et al., 2013), with accompanying heat maps generated using GraphPad Prism 7.0.

### Immunoblotting

Expression levels of Cno transgenic proteins and knockdown efficiency of *Rap1* and *cno* were determined by Western blotting. Embryo lysates were prepared by grinding dechorionated embryos in ice-cold lysis buffer (1% NP-40, 0.5% Na deoxycholate, 0.1% SDS, 50 mM Tris pH 8, 300 mM NaCl, 1.0 mM DTT and Halt^TM^ protease and phosphatase inhibitor cocktail [Thermo Scientific, Rockford, IL]). Lysates were cleared at 16,000 x g and protein concentration determined using the Bio-Rad Protein Assay Dye (Bio-Rad, Hercules, CA). Lysates were resolved by 8 or 12% SDS-PAGE, transferred to nitrocellulose filters and blocked for 1 h in 10% dry milk powder in PBS-Tw (0.1% Tween-20 in PBS). Membranes were incubated in primary antibody for 2 h (see Table 1 for antibody concentrations); washed four times for 5 min each in PBS-Tw; and incubated with IRDye-coupled secondary antibodies for 45 min. Signal was detected over a 4-log dynamic range with the Odyssey CLx infrared imaging system (LI-COR Biosciences, Lincoln, NE). Band densitometry was performed using ImageStudio software version 4.0.21 (LI-COR).

## Author contributions

T.T. Bonello, K.Z Perez-Vale, and M. Peifer conceived the study. K.D. Sumigray generated the transgenic wildtype and mutant *cno* transgenes, and K.Z Perez-Vale analyzed *dizzy* mutants and helped analyze the role of the RA-domains. All other experiments were carried out by T.T. Bonello. T.T Bonello and M. Peifer wrote the manuscript with input from the other authors.

## Acknowledgements

Thanks to W. Choi for developing the anti-Rap1 and anti-Baz antibodies. Thanks to R. Reuter for key Drosophila stocks, M. Price for advice in 3D imaging, Peifer lab members including A. Spracklen for helpful advice and comments, and J. Grosshans for helpful discussions. This work was supported by NIH R35 GM118096 to M.P. T.T.B. was supported in part by a Sir Keith Murdoch Fellowship from the American Australian Association. K.Z.P-V. was supported by NIH T32 GM007092 and NIH R25 GM055336. K.D.S was supported by NIH F32 GM106516

---

**Supplementary Figure 1.**
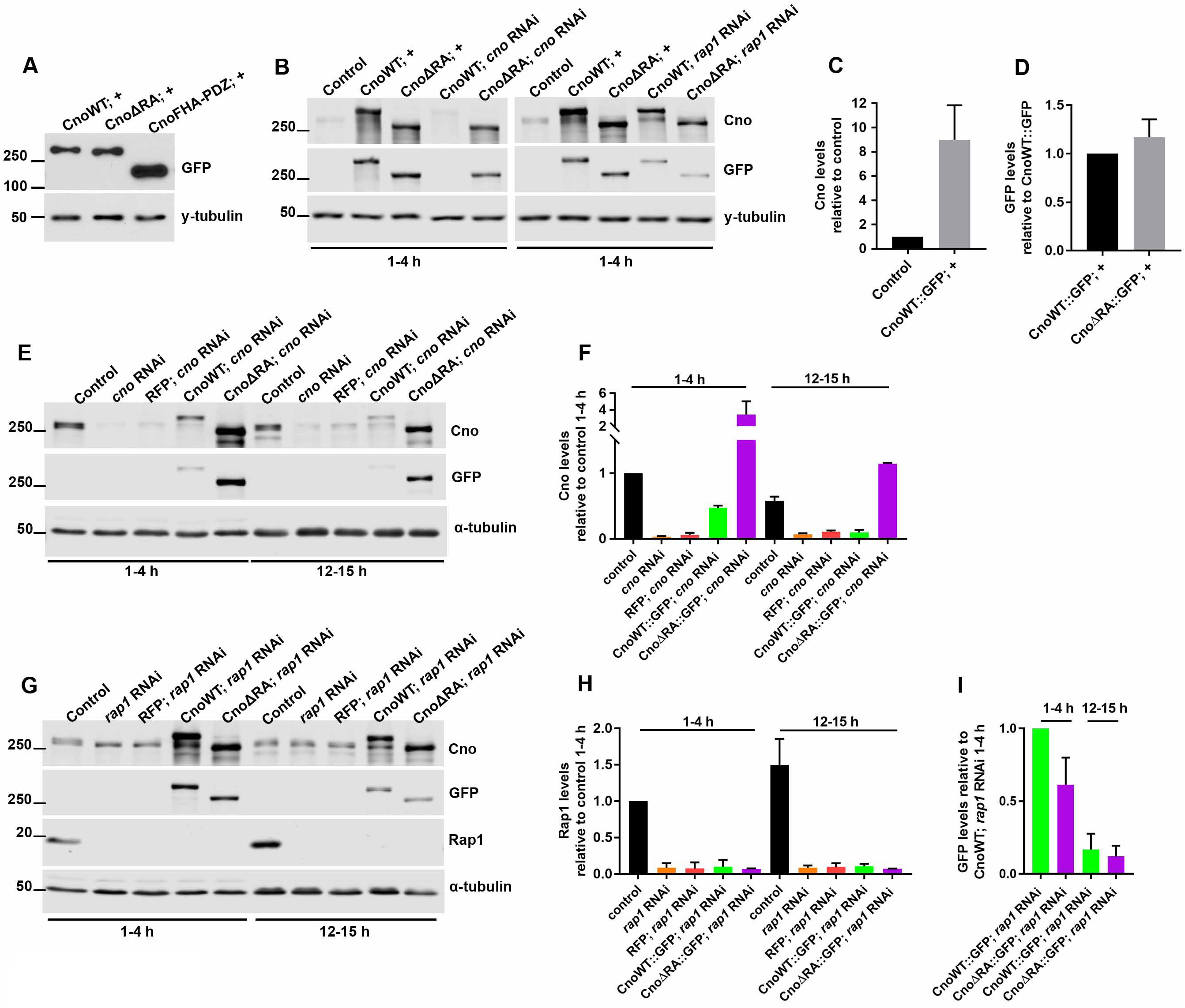
Effectiveness of *cno* and *Rap1* RNAi knockdown, and expression levels of CnoWT or CnoΔRA in wildtype, Cno or Rap1-depleted backgrounds. (A,B) Western blots showing levels of GFP-tagged Cno transgenes relative to endogenous Cno. (C) CnoWT::GFP is expressed 9-fold over endogenous Cno levels. (D) CnoWT::GFP and CnoΔRA::GFP are expressed at equivalent levels in wildtype background. (E) Western blot showing expression of GFP-tagged Cno transgenes in a *cno* RNAi background and (F) corresponding quantification of Cno levels. (G) Western blot showing expression of GFP-tagged Cno transgenes in *Rap1* RNAi background. (H) Rap1 levels are highly reduced in early embryo collections (1–4 h) and levels do not return by late embryogenesis (12–15 h). (I) CnoWT::GFP and CnoΔRA::GFP are expressed at relative levels in *Rap1* RNAi background.

**Supplemental Figure 2.**
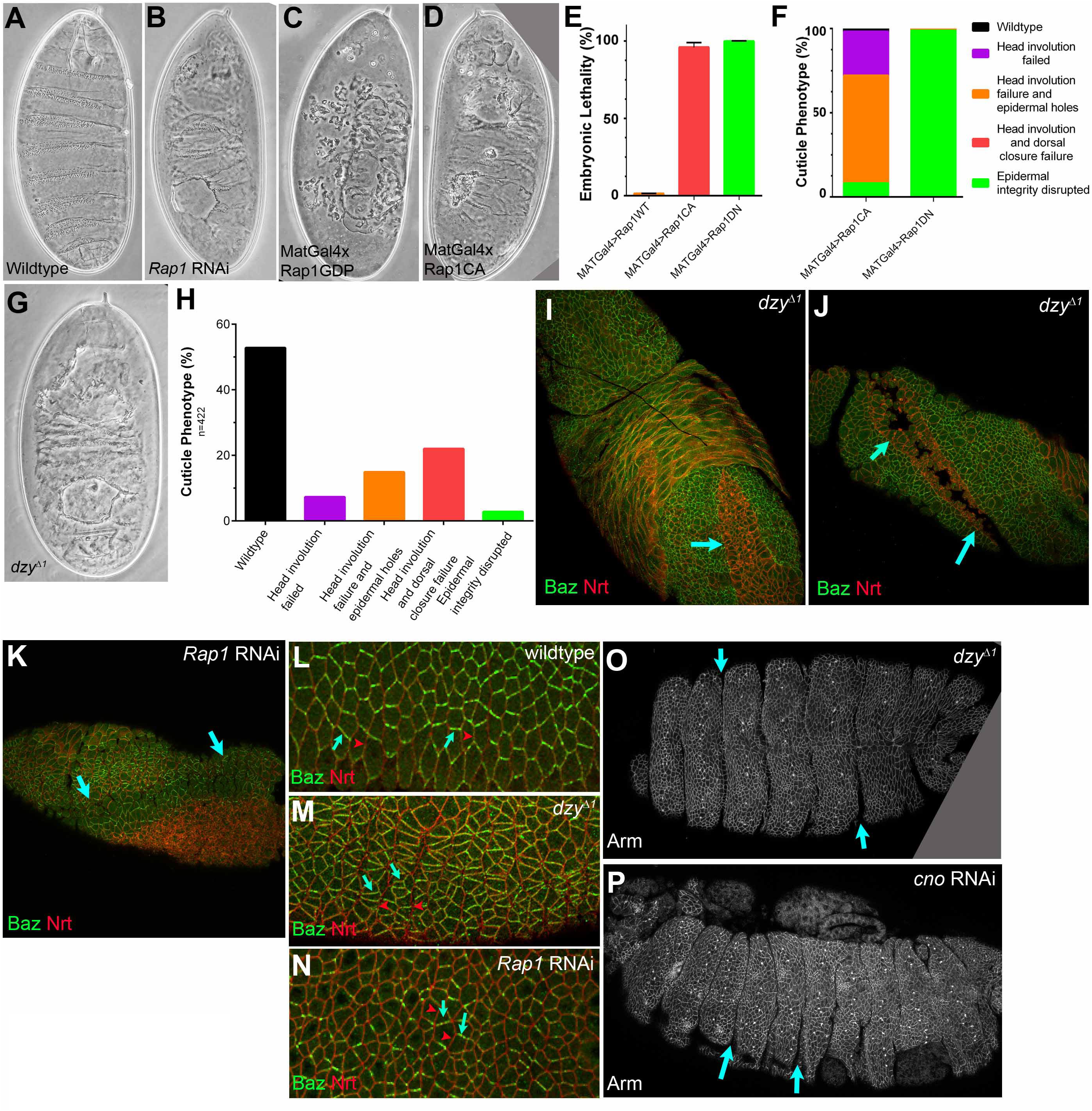
Phenotypes of Rap1 activity mutants and the Dizzy Rap1 GEF. (A-D, G) Cuticle preps. (A) Wildtype. (B) *Rap1* RNAi leads to defects in head involution and dorsal closure and reduces epidermal integrity. (C) Expressing Rap1GDP leads to strong disruption of epidermal integrity. (D) Expressing Rap1CA leads to defects in head involution and dorsal closure with modest effects on epidermal integrity. (E) Both Rap1 activity mutants lead to highly penetrant embryonic lethality, while expressing Rap1WT has no effect. (F) Quantification of effects of Rap1 activity mutants on morphogenesis, as assessed by cuticle analysis. (G) Maternal/zygotic *dzy* mutants have defects in head involution and dorsal closure and reduced epidermal integrity. (H) Quantification of effects of *dzy* mutants on morphogenesis, as assessed by cuticle analysis. The fraction with near wildtype cuticles include the 50% of the embryos who are zygotically-rescued. (I) *dzy* mutants fail to fully internalize their mesoderm (arrow=open ventral furrow; Spahn et al., 2012), thus resembling *cno* and *Rap1* mutants (Sawyer et al., 2009). (J, K) *dzy* mutants also have a twisted germband (J, arrows), suggesting a failure in germband extension. This is more severe than the delayed germband extension seen in *cno* mutants (Sawyer et al., 2011), but we observed a similar twisted germband phenotype after *Rap1* RNAi (K, arrows). (L-N) Lateral ectoderm, stage 8. (L) In wildtype, Baz is planar-polarized, accumulating more on dorsal-ventral (arrows) than on anterior-posterior (arrowheads) borders (Zallen and Wieschaus, 2004). (M) This planar polarity is dramatically enhanced in *dzy* mutants, with Baz lost from anterior-posterior borders (arrowheads). This mimics the effect of *cno* mutants (Sawyer et al., 2011). (N) *Rap1* RNAi embryos have a similar enhancement of Baz planar polarity, though Baz staining is already becoming more fragmented, perhaps presaging loss of epithelial integrity. (O, P) Stage 14 *dzy* mutants have abnormally deep segmental grooves (arrows), a phenotype also seen in *cno* zygotic mutants (Choi et al., 2011) or after *cno* RNAi (P).

**Supplementary Figure 3.**
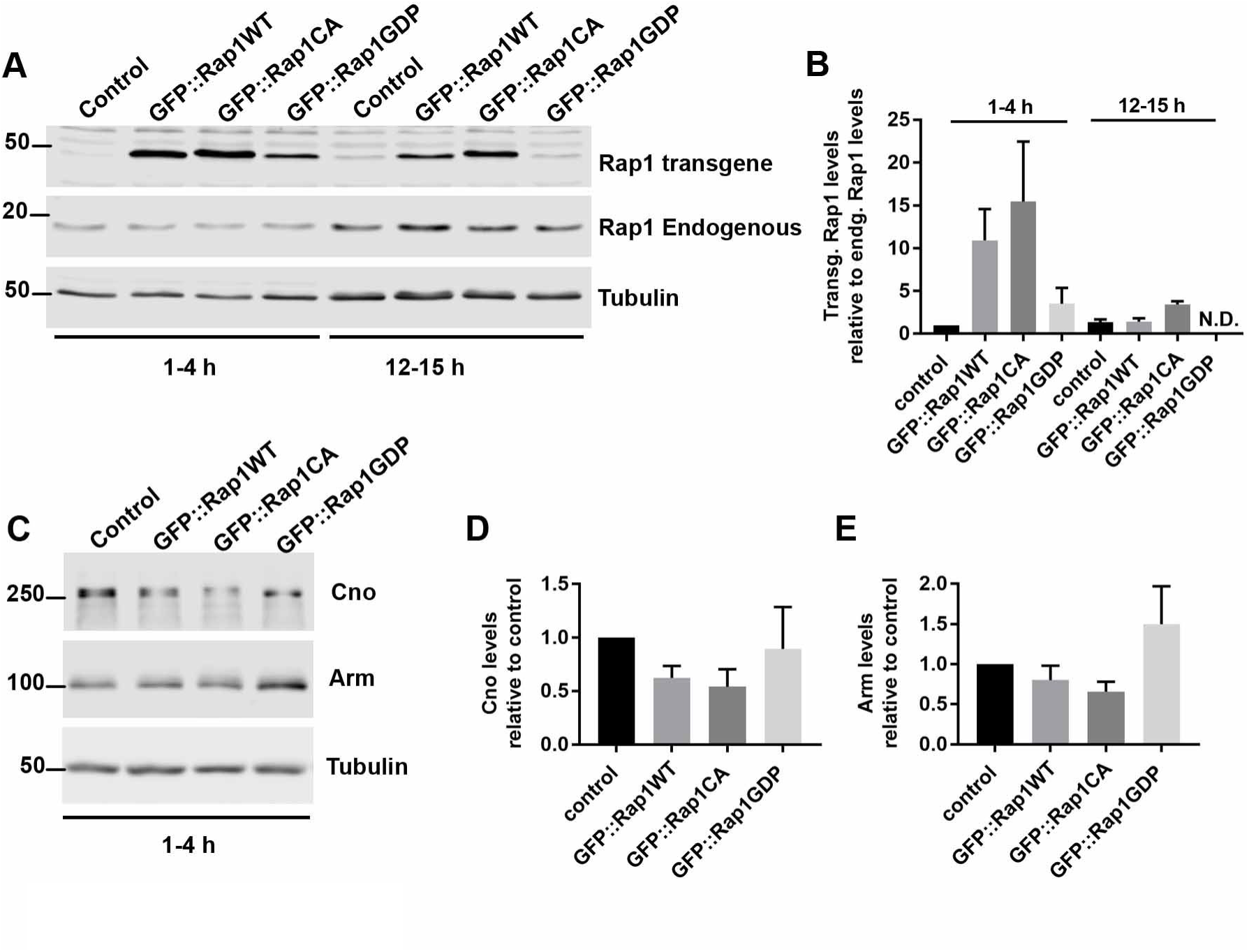
Absolute levels of Rap1 activity mutants relative to endogenous Rap1. (A) Western blot and (B) corresponding quantification showing expression levels of Rap1 activity mutant protein relative to endogenous Rap1. Rap1GDP is not detectable (N.D.) in 12–15 h embryos. (C) Western blot and corresponding quantification of (D) Cno and (E) Arm levels in Rap1 activity mutants.

